# A latent tau amplifier in the neuronal proteome lowers the seeding threshold for Alzheimer’s disease pathology

**DOI:** 10.64898/2026.11.16.766884

**Authors:** Grigoria Tsaka, Rajasekhar Kolla, Anne E. Wiedmer, Alexandros Georgilis, Sara Kenis, Gerd Van der Hoeven, Filip Claes, Ladan Khodaparast, Laleh Khodaparast, Bas Lahaije, Sandra O. Tomé, Kevin Muwonge, Ramon Duran, Rik Vandenberghe, Christine A.F. von Arnim, Nikolaos Louros, Dietmar R. Thal, Frederic Rousseau, Joost Schymkowitz

## Abstract

The stereotyped spread of tau pathology through connected brain regions is accompanied by striking differences in the vulnerability of individual neuronal populations, pointing to cell-intrinsic determinants of susceptibility. Using a combination of computational APR analysis, in vitro and cellular aggregation assays, and quantitative proteomics of human postmortem brain, we identify the splicing factor TRA2B as an endogenous modifier of this susceptibility. TRA2B enhances the activity of patient-derived tau seeds, promotes tau pathology in vivo, and progressively redistributes from the nucleus into insoluble tau-associated material in Alzheimer’s disease brain. Mechanistically, TRA2B contains two aggregation-prone regions (APRs) homologous to tau nucleation sites that are required for tau interaction and seeding enhancement. Yet this activity is not constitutive: isolated TRA2B APRs promote aggregation, whereas the full-length protein can inhibit, amplify or have little effect on tau aggregation depending on protein state and seeding context. Perturbation of the regulatory RS2 region switches TRA2B away from its amplifying state, revealing that its sequence-encoded aggregation potential is regulatable. Thus, TRA2B acts as a latent tau amplifier, providing a mechanism by which the neuronal proteome can tune cellular susceptibility to the propagation of pathological tau.

## INTRODUCTION

Tau pathology in Alzheimer’s disease (AD) and related tauopathies spreads through the brain in a stereotyped pattern (1, 2). Misfolded tau propagates from neuron to neuron by a prion-like seeding mechanism, templating the conformational conversion of soluble tau monomers and progressively recruiting new regions into the advancing front of pathology (3–8). This process is neither inevitable nor uniform: patient-derived tau preparations span orders of magnitude in seeding potency, many tau-containing neurons never develop overt neurofibrillary tangles, and the rate at which pathology spreads varies between individuals (9–11). One variable underlying this heterogeneity is the seeding threshold: the minimum aggregate potency required for productive tau propagation. This threshold has received little mechanistic attention (12), yet the cellular factors that modulate it bear directly on why tau pathology is regionally selective, why some individuals progress rapidly while others plateau, and why tau-directed therapies have proven difficult to translate (9).

Among the cellular factors likely to influence tau seeding behaviour, RNA-binding proteins (RBPs) are prominent candidates (13–15). TIA-1, HNRNPs, MSI1/2, G3BP2 and TDP-43 are among the most consistently identified tau-associated proteins across independent human AD interactome and proteome datasets, indicating that RBPs are functional participants in tau pathology rather than incidental bystanders (13, 16). For individual proteins the evidence is substantial: TIA-1 converts tau into neurotoxic oligomers through RNA-mediated phase separation (17); G3BP2 maintains tau solubility, and its loss worsens tau pathology (18); and TDP-43, whose nuclear clearance and cytoplasmic accumulation co-occur with tau in multiple tauopathies, engages tau aggregates through mechanisms that remain incompletely defined (19–21). What has been missing is a molecular principle for predicting which RBP interactions actively drive tau aggregation rather than reflecting co-sequestration into inclusions.

Aggregation-prone regions (APRs), short hydrophobic segments that nucleate protein misfolding through β-strand stacking, provide a basis for this distinction (22–24). Proteins whose primary sequences encode APRs homologous to those of tau can co-aggregate with tau in a sequence-specific manner, modifying its nucleation kinetics, fibril morphology and cellular spreading behaviour (25). The interaction is not promiscuous: its direction and strength are determined by the structural compatibility between the homologous APR and the tau amyloid core, and disrupting the APR sequence abolishes it. By scanning the human proteome for PHF6-homologous sequences and testing full-length candidates in a tau biosensor cell model, we previously identified TRA2B as one of the most potent tau aggregation enhancers in this class, capable of sensitising cells to tau seeding at picomolar seed concentrations (25). The mechanism and disease relevance of this threshold-lowering effect were not established.

TRA2B is a 288-amino acid RNA splicing factor with established links to tau biology. It controls alternative splicing of the tau pre-mRNA and is recurrently detected in proteomic analyses of neurofibrillary tangles and tau-enriched fractions from AD brain, placing it at the intersection of splicing regulation and tau pathology (13, 26, 27). Re-examination of the TRA2B sequence reveals that it encodes two APRs homologous to the major aggregation nucleation sites of tau: APR1 is homologous to the PAM4 motif (23), and APR2 to the PHF6 motif (VQIVYK), a tau APR characterised after our initial screen was performed (28). Both APRs are are predicted to be surface-exposed within the RNA recognition motif (RRM) of TRA2B. This arrangement predicts cooperative, multivalent engagement of both tau nucleation sites, and therefore a stronger interaction than the single-contact model previously assumed. Here we establish the disease relevance and molecular mechanism of the threshold-lowering activity of TRA2B. We show that it operates on patient-derived tau seeds across the range of disease-relevant potencies, amplifies tau seeding competence in vivo, and is accompanied by a redistribution of TRA2B in AD brain. The mechanism we describe, a seed-gated switch that keeps the co-aggregating activity of TRA2B dormant until a seeding template is present, provides a framework for how the normal proteome can harbour latent amplifiers of tau pathology.

## RESULTS

### TRA2B enhances tau seeding

The seeding potency of tau extracted from AD brain varies considerably between patients, reflecting differences in aggregate load, post-translational modifications and structural strain composition. We reasoned that if TRA2B lowers the seeding threshold, its effects might be most pronounced on weak seeds, close to or below the threshold for productive propagation. We therefore established individual dose-response seeding curves for sarkosyl-insoluble tau from eight AD brain cases spanning a range of Braak stages, using HEK293 tau biosensor cells (Fig. 1A). For each case, we identified a seed dilution within the dynamic range of the assay, avoiding both saturation and absence of signal.

**Figure 1.**
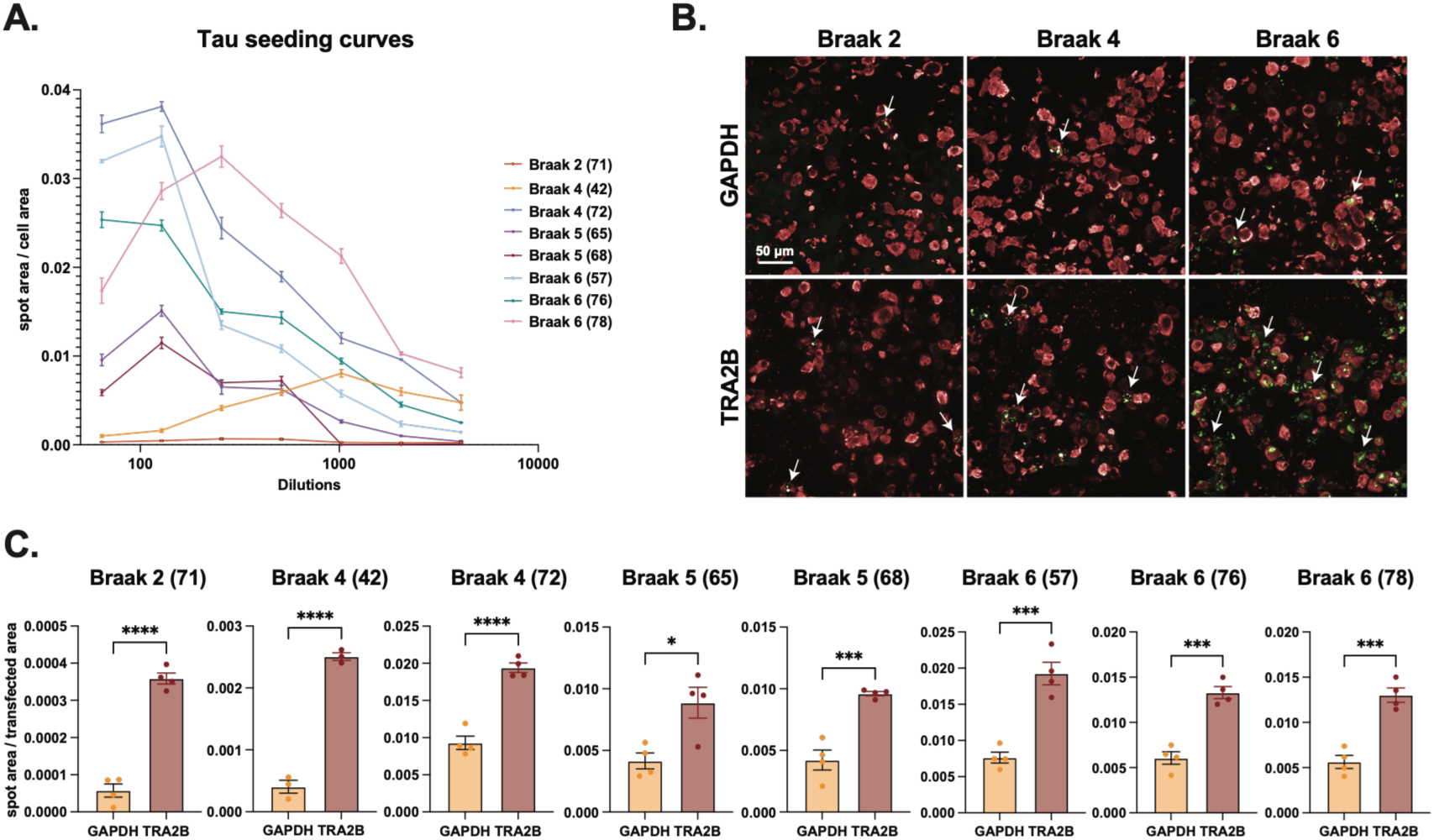
TRA2B inverts the propagation fate of sub-threshold tau seeds. **A)** Individual dose-response seeding curves for sarkosyl-insoluble tau from eight AD patient brains (cases 71, 42, 72, 65, 68, 57, 76, 78; **Table 1**), measured in tau biosensor cells. Curves span multiple Braak stages and establish the dynamic range and working dilution for each case. **B)** Representative fluorescence images of tau biosensor cells co-transfected with calibrated patient-derived tau preparations and either GAPDH (control) or TRA2B. FRET-positive puncta indicate tau aggregates. Scale bar applies to all images. **C)** Quantification of FRET-positive puncta across all eight cases. TRA2B significantly enhances seeding across all samples, but the fold-change is inversely correlated with baseline seeding activity: cases 71 and 42, with the lowest intrinsic potency, show the largest responses, consistent with threshold-lowering rather than uniform enhancement (n = 4; unpaired t-test).

Across all eight cases taken forward, TRA2B expression increased FRET-positive tau aggregate formation relative to GAPDH controls, although the magnitude of the effect varied between cases (Fig. 1B–C). Cases 71 and 42, which showed the lowest intrinsic seeding activity in their dose-response profiles, gave the largest fold changes upon TRA2B expression. At the tested dilution, these preparations produced little aggregation in control cells but became robustly seeding-competent in TRA2B-expressing cells. The effect of TRA2B was inversely related to baseline seed potency, with the largest relative enhancement observed for the weakest seeds. These findings are consistent with TRA2B lowering the threshold for productive tau seeding.

### TRA2B drives accumulation of seeding-competent tau in the living brain

PS19 transgenic mice express P301S human tau at approximately fivefold above endogenous murine tau levels and develop progressive tau pathology with age (12, 29). To define the temporal progression of tau pathology in the PS19 model and establish the sensitivity window of our experimental readouts, we first conducted a longitudinal characterisation across both sexes spanning ages from 2 to 10 months. A multi-modal approach was employed, integrating functional seeding assays using sarkosyl-insoluble tau fractions, immunohistochemical evaluation of tau pathology (pS202/T205 tau, AT8), and quantitative assessment of phosphorylated tau species by ELISA (pS199 tau). Tau seeding competence emerged at an early stage, with detectable seeding activity observed as early as 2 months of age, significantly preceding overt histopathological changes (**Suppl. Fig. 1A-B**). Seeding potency increased progressively with age, indicating an accumulation of aggregation-competent tau species over time. In contrast, biochemical detection of pathological phosphorylated tau by ELISA became apparent at approximately 4 months (**Suppl. Fig. 1C)**, while immunohistochemical detection of tau pathology was first observed at 5 months, initially at low levels (**Suppl. Fig. 1A-B**). A marked escalation in pretangle formation and neurofibrillary tangle deposition occurred after 8 months. Tau seeding was therefore detectable before the appearance of conventional histopathological markers. Importantly, this temporal sequence is consistent with previously reported characterisations of the PS19 model (12, 29).

Guided by this characterisation, we next asked whether TRA2B’s threshold-lowering effect on patient-derived seeds translates into a disease-relevant phenotype in vivo. We delivered AAV-encoded human TRA2B or vehicle (PBS) unilaterally into the hippocampus of two-month-old PS19 mice, before overt pathology, and harvested tissue at six months, capturing four months of TRA2B overexpression across the period of tau aggregation onset and early spread. Sarkosyl-insoluble tau from TRA2B-overexpressing hippocampi generated substantially elevated seeding activity in tau biosensor cells relative to PBS-injected controls, with seeding curves shifted upward across dilutions and a significant increase in FRET-positive puncta at normalised concentrations (**Fig. 2A-C**). Total tau measured by ELISA was also higher in TRA2B-overexpressing animals (**Fig. 2D**). TRA2B overexpression therefore increased both total tau abundance and the seeding activity of the insoluble fraction. These findings indicate that TRA2B can enhance the accumulation of seeding-competent tau in vivo.

**Figure 2.**
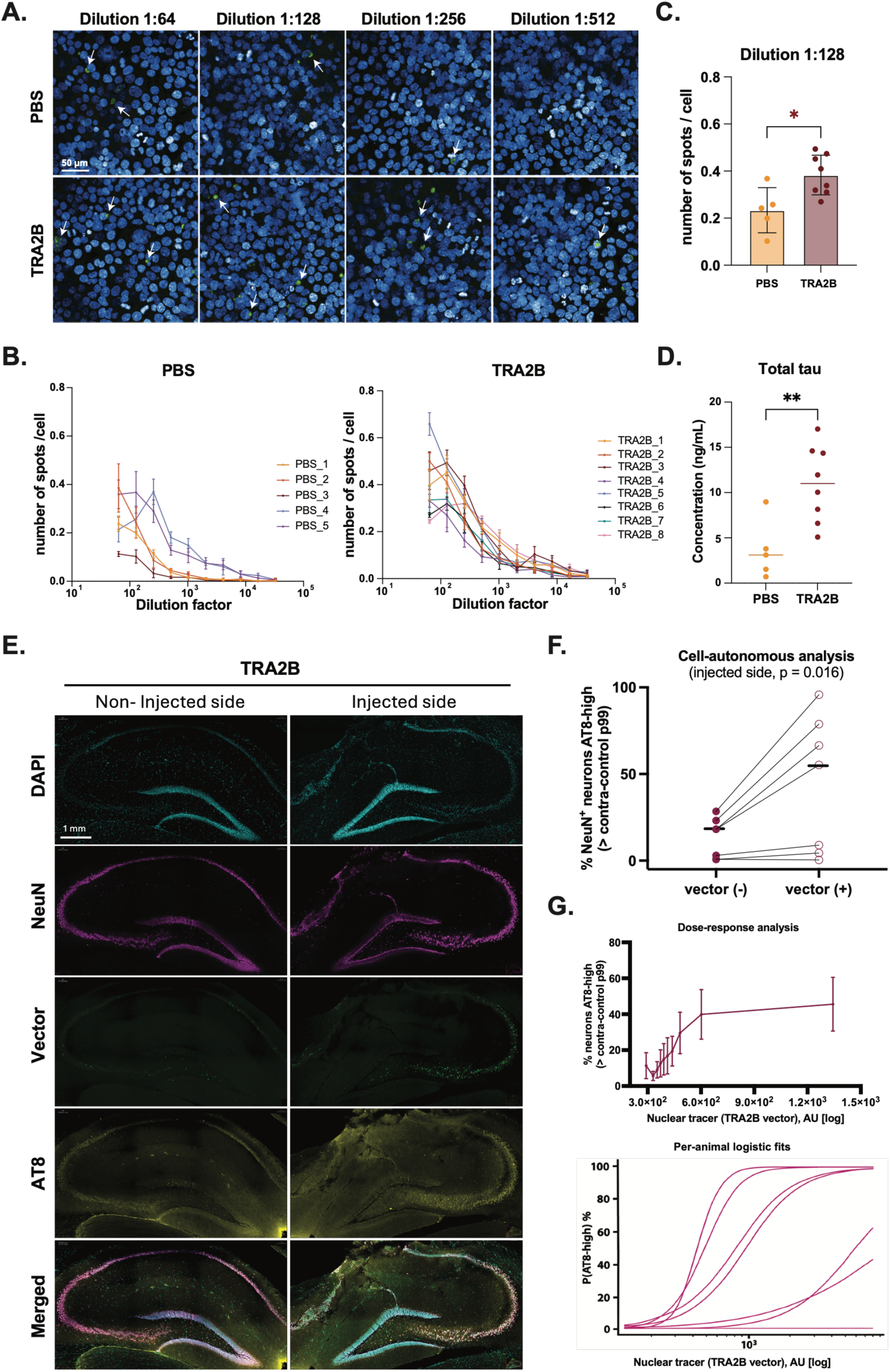
TRA2B overexpression expands the pool of seeding-competent tau in vivo. **A)** Representative images of tau biosensor cells treated with sarkosyl-insoluble fractions from PS19 mice injected with vehicle (PBS) or AAV-TRA2B. Scale bar applies to all panels. **B)** Seeding curves shifted upward in TRA2B-overexpressing animals across all dilutions tested (n = 4 per group). **C)** Quantification at normalized dilution (1:128), showing significantly elevated seeding activity in AAV-TRA2B mice, reflecting a higher burden of seeding-competent tau species in vivo (unpaired t-test, n = 5-8). **D)** Total tau ELISA of hippocampal homogenates: TRA2B overexpression drives both tau accumulation and enrichment of seeding-competent species (unpaired t-test, n = 5-8). **E)** Representative images of TRA2B-injected and contralateral non-injected hippocampi immunostained for DAPI (nuclei), NeuN (neurons), HA (TRA2B vector expression), and AT8 (pS202/T205 tau). **F)** Cell-autonomous association of TRA2B expression with pathological tau. For each TRA2B animal (n = 7), the percentage of injected-hippocampus NeuN⁺ neurons that are AT8-high (> contralateral Vector⁻ control 99^th^ percentile), among vector-expressing (Vector⁺) versus non-expressing (Vector⁻) neurons of the same section. Lines pair the two populations within an animal; black bars, medians. One-sided Wilcoxon signed-rank (Vector⁺ > Vector⁻). **G)** Dose-dependence. Top: percentage of neurons that are AT8-high as a function of nuclear TRA2B-vector (Tracer) level, binned into deciles; mean ± SEM across animals. Bottom: per-animal logistic fits of P(AT8-high) on nuclear vector intensity; slope positive in 7/7 animals (one-sided Wilcoxon p = 0.008).

### TRA2B-expressing neurons accumulate pathological tau cell-autonomously, at an early disease stage

To test whether the increased tau seeding and total tau measured in TRA2B-injected PS19 mice arise in the neurons that actually express TRA2B, we performed single-cell quantification of phosphorylated tau (AT8), the neuronal marker NeuN, and the AAV-TRA2B vector (“Tracer”, nuclear) in the same hippocampal sections, classified cell-by-cell (**Fig. 2E**). At this early age the two groups were not separable at the gross level: the fraction of AT8-positive neurons did not differ significantly between TRA2B- and PBS-injected animals (injected hemisphere; Mann-Whitney P > 0.3) (**Suppl. Fig. 2A-B**), and, reflecting the injection procedure itself, an injected-versus-contralateral enrichment of AT8-positive neurons was also present in PBS controls (**Suppl. Fig. 2A, C**). Detecting the TRA2B-specific effect therefore required resolving tau pathology cell-by-cell against the neurons that actually express the vector.

Within the injected hippocampus, neurons expressing the TRA2B vector (Vector⁺) were AT8-high far more frequently than non-expressing (Vector⁻) neurons in the same section (**Fig. 2F**). Across animals the AT8-high fraction was higher in Vector⁺ neurons in 6 of 7 mice (median 55.2% vs 18.1% of neurons above the control 99th-percentile AT8 gate; median odds ratio ≈ 6; paired one-sided Wilcoxon signed-rank P = 0.016; the same at the 95th-percentile gate). Because this comparison is made between neighbouring neurons on the same physical section, it is independent of staining or imaging variation and of any spread of vector to the contralateral hemisphere.

The effect was dose-dependent rather than all-or-none: the probability that a neuron was AT8-high rose monotonically with its nuclear TRA2B-vector level, from ∼6% of neurons in the lowest expression decile to ∼46% in the highest (**Fig. 2G**). A per-animal logistic regression of AT8-high status on nuclear vector intensity gave a positive slope in all 7 animals (one-sided Wilcoxon P = 0.008). The single mouse without a clear Vector⁺ vs Vector⁻ difference had very low tau pathology overall, so little tau was available to partition.

At the hemisphere level, the injected-side AT8-high enrichment was significant within the TRA2B group (**Suppl. Fig. 2C**). We treat this as supporting rather than primary evidence, however, because the same injected-versus-contralateral enrichment is seen in PBS controls (it reflects the injection itself) and because it is further blunted by spread of vector to the contralateral hemisphere (a minority of contralateral neurons, typically 0.4–2%, versus 4–23% on the injected side, express the vector). One animal showed substantial bilateral transduction (about 40% of contralateral neurons vector-positive) and was excluded from the hemisphere-level comparison; its injected hemisphere was retained for the cell-autonomous analysis, where the comparison is made between vector-positive and vector-negative neurons of the same hemisphere and is therefore unaffected by the injection or by vector spread.

Together, at a disease stage that precedes overt tangle pathology, TRA2B overexpression increases pathological, AT8-positive tau specifically in the neurons that express it, in a dose-dependent manner. This single-cell, cell-autonomous phenotype is directionally concordant with the bulk increases in seeding-competent and total tau measured in the same cohort (**Fig. 2B-D**), linking TRA2B expression to tau pathology at the level of the individual neuron.

### TRA2B undergoes pathological sequestration into tau aggregates in AD brain

We performed quantitative proteomic analysis of postmortem brain tissue from two regions with distinct temporal involvement in AD, perirhinal cortex (BA36, early-affected) and frontal cortex (BA6, late-affected), sequentially fractionated into a TBS-insoluble but dispersible fraction (DISP) and an SDS-soluble fraction. These fractions report on different assembly states: the DISP fraction contains loosely associated, largely oligomeric material that remains extractable without detergent, whereas the SDS fraction contains more tightly packed assemblies that are released only by detergent. A shift of a protein between the two therefore reflects its incorporation into more stable, higher-order aggregates. TRA2B showed a disease-stage-dependent change in solubility. (**Fig. 3A-C**). In BA36, TRA2B was unchanged across non-demented controls, asymptomatic AD and symptomatic AD, consistent with the relative preservation of this region in early disease. In BA6, a reciprocal shift emerged in symptomatic AD: TRA2B was depleted from the DISP fraction while accumulating in the SDS fraction, a redistribution confirmed by independent ELISA quantification in a validated subcohort (**Fig. 3C**).

**Figure 3.**
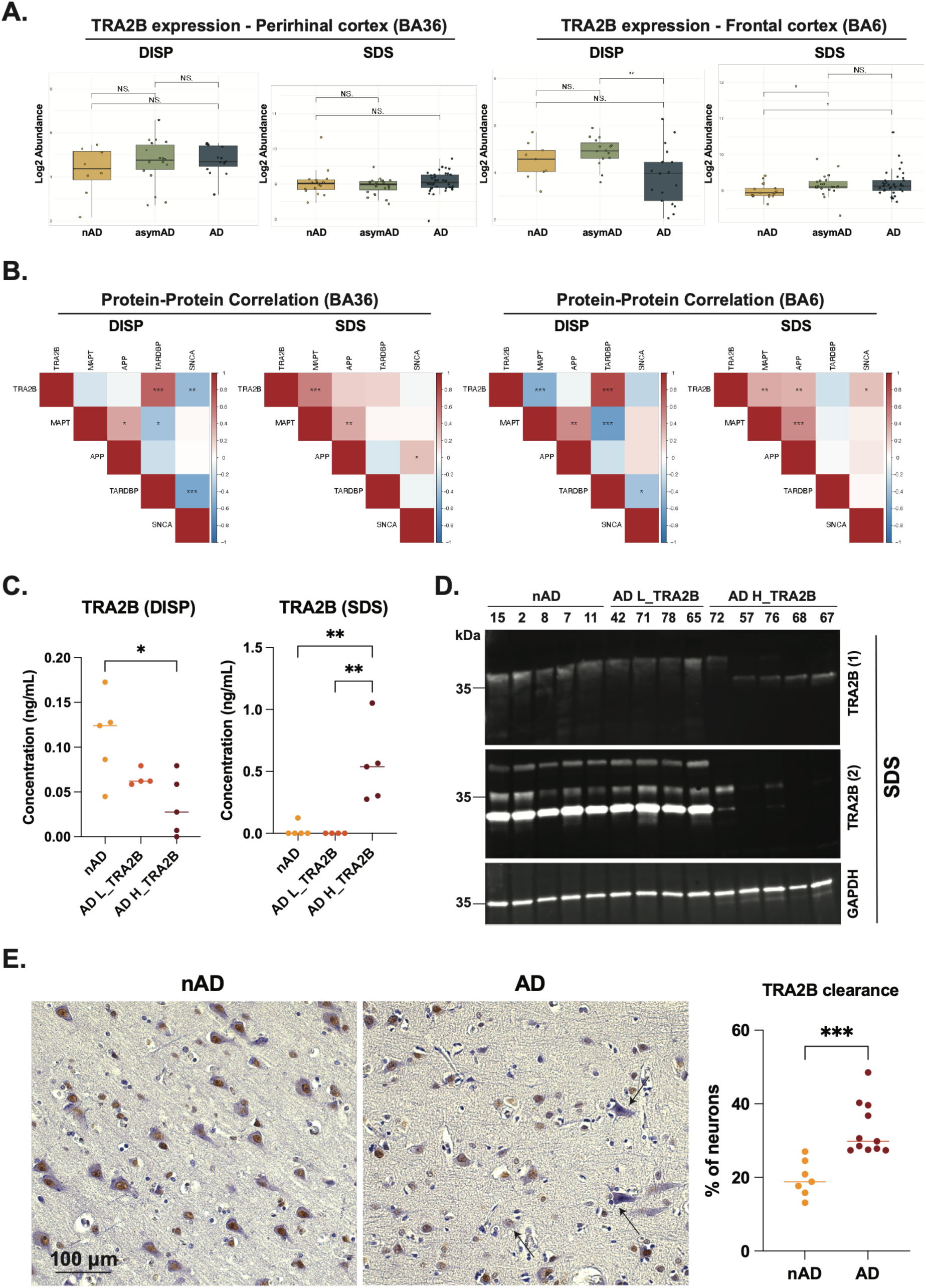
TRA2B is progressively sequestered into tau pathology in AD brain. **A)** Proteomic quantification of TRA2B in TBS-insoluble dispersible (DISP) and SDS-soluble fractions of perirhinal cortex (BA36) and frontal cortex (BA6) from non-demented controls (nAD), asymptomatic AD (asymAD), and symptomatic AD. In late-affected BA6, symptomatic AD cases show a reciprocal shift, with TRA2B depleted from the DISP fraction and enriched in the detergent-resistant SDS fraction, consistent with its recruitment into more stable, higher-order assemblies. **B)** Protein co-expression correlation of TRA2B with tau (MAPT), APP, TDP-43 (TARDBP) and alpha-synuclein (SNCA). In the DISP fraction, TRA2B correlates with TDP-43 (nuclear function); in the SDS fraction it instead correlates with tau, marking this detergent-resistant pool as the disease-associated one. The rewiring of TRA2B’s co-expression network between fractions places its aggregated pool in a distinct, tau-linked protein environment. **C)** ELISA validation of the solubility shift in a validated BA6 subcohort stratified as nAD, AD L_TRA2B, and AD H_TRA2B (one-way ANOVA, n = 4-5). **D.** Immunoblot of the SDS fraction (BA6): multiple anti-TRA2B antibodies fail to detect, or detect only a modified form of, TRA2B in AD H_TRA2B cases, indicating conformational remodeling or incorporation into higher-order assemblies. GAPDH loading control. **E)** Quantification of TRA2B nuclear clearance in BA6 neurons (cases 1, 7, 8, 11, 12, 16, 42, 53, 55, 56, 57, 64, 65, 67, 68, 72, 78, 81): advanced AD cases show a significantly higher proportion of TRA2B-depleted nuclei than controls, establishing that tau sequestration depletes the neuronal nuclear TRA2B pool (unpaired t-test, nAD n = 7, AD n = 11).

The two compartments differ in their co-expression signatures. In the DISP fraction, TRA2B abundance correlated strongly and specifically with TDP-43 across both brain regions, consistent with shared nuclear residency and RNA-processing function (**Fig. 3B**). This relationship was absent in the more aggregation-rich SDS fraction, replaced by a positive correlation with tau. In late-affected BA6, SDS-soluble TRA2B additionally co-varied with Aβ and α-synuclein, suggesting incorporation into a broader aggregation-associated proteome. The proteins associated with TRA2B abundance therefore differed between the two fractions.

The TRA2B species that accumulates in the SDS fraction of symptomatic AD cases is biochemically altered. Multiple anti-TRA2B antibodies targeting distinct epitopes either fail to detect TRA2B in the SDS fraction of high-TRA2B AD cases, or detect only a modified form, a pattern consistent with conformational remodelling, truncation or steric shielding within a higher-order assembly (**Fig. 3D, Suppl. Fig. 3**). The most parsimonious interpretation is that TRA2B has been co-incorporated into insoluble tau-containing structures in a manner that modifies its antibody-accessible epitopes.

At the cellular level, this sequestration manifests as neuronal nuclear clearance: neurons in the frontal cortex of advanced AD cases show a significantly higher proportion of TRA2B-depleted nuclei than non-demented controls (**Fig. 3E**). The pattern parallels TDP-43 pathology, but the biochemical association with tau suggests a distinct basis: tau-specific co-aggregation driven by sequence-encoded APR interactions rather than generic RBP mislocalisation. Taken together, these data indicate that a nuclear splicing factor, which is also the regulator of the alternative splicing of tau itself, is progressively depleted from the nucleus and incorporated into tau aggregates as disease advances.

### TRA2B encodes dual sequence homology to both major tau aggregation nucleation sites

The molecular basis for the TRA2B-tau interaction lies in two APRs within its RNA recognition motif. APR1 (VFSKYG) is homologous to the tau PAM4 motif (VQSKIG), a more recently characterised aggregation-prone region that contributes to fibril stability across multiple tau strains. APR2 (VSIVYD) is homologous to the tau PHF6 motif (VQIVYK), the best-characterised tau nucleation site and a stabiliser of all known tau amyloid polymorphs (**Fig. 4A**). Both APRs are surface-exposed within the structured RRM of TRA2B and are separated by only four residues, an arrangement that could place them within the spatial register of the PHF6 and PAM4 stacking interfaces simultaneously when TRA2B engages a growing tau fibril.

**Figure 4.**
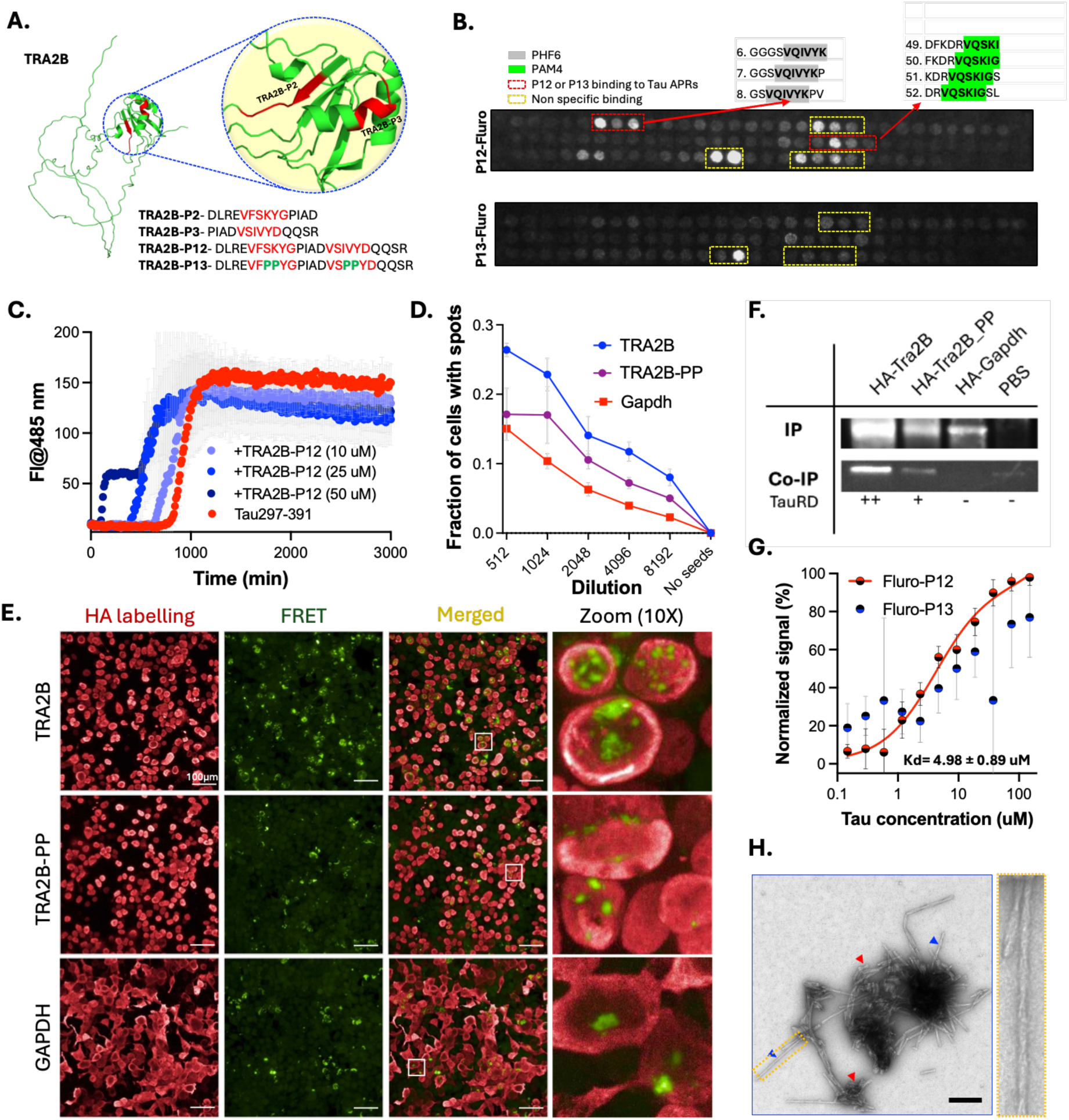
*In vitro* aggregation of Tau and TRA2B-P12. **A)** AlphaFold-predicted structure of TRA2B highlighting the localization of aggregation-prone regions (APRs) within the structured region of the protein. Sequences of the TRA2B-derived peptides used in the in vitro experiments are shown. **B)** Peptide array mapping of TRA2B-P12 binding on tau. Overlapping tau peptides were synthesized on a cellulose membrane (SPOT synthesis) and probed with fluorescently labelled TRA2B-P12 and TRA2B-P13. **C)** Thioflavin T (ThT) aggregation kinetics of Tau297-391 in the presence of increasing concentrations of TRA2B-P12 (10, 25, and 50 μM). **D)** Seeding curves of Tau aggregates following dose-dependent treatment with human Tau seeds in biosensor cells overexpressing wild-type TRA2B, proline-substituted mutant TRA2B-PP, or GAPDH (control). Bar plot (right) highlights differences observed at the highest seed concentration. Data are represented as mean values ± SD (n = 3 independent experiments). **E)** Representative images of human Tau seeding (FRET, green) in biosensor cells overexpressing TRA2B, TRA2B-PP, or GAPDH (HA-labelling, red). Scale bar = 100 μm. **F)** Co-Immunoprecipitation of TauRD with HA-tagged TRA2B, TRA2B-PP, or GAPDH. Upper panel shows immunoprecipitated HA-tagged constructs (IP); lower panel shows co-immunoprecipitated TauRD (Co-IP). **G)** Binding of Fluor-P12 and Fluor-P13 to Tau297-391 (Fluor-P12, Red trace, 4.98 ± 0.89 uM). No reliable binding constant was observed for Fluor-P13. Curves are shown as mean values ± SD (n = 4 or 3). **H)** TEM image of fibrillar aggregates from Tau297-391 and TRA2B-P12 after 72 h. Red arrowheads indicate short fibrils resembling TRA2B-P12 or Tau fibrils, whereas blue arrowheads indicate long, singular twisted fibrils. Scale bar for TEM image, 200 nm.

We tested the sequence specificity of this interaction by peptide array. Fluorescein-labelled TRA2B-P12 (a 24-residue peptide encompassing both APRs) and TRA2B-P13 (a proline-substituted variant in which both APRs are disrupted) were applied to overlapping 10-residue tau peptide arrays spanning Tau297-391 (**Fig. 4A-B**). TRA2B-P12 produced binding signals at the PHF6 and PAM4 regions of the tau sequence, whereas TRA2B-P13 showed markedly reduced or absent signal at these positions, implicating the APRs in sequence-specific recognition. Off-target binding was also detected for TRA2B-P12; because these signals were retained in TRA2B-P13, they originate from the flanking regions of the peptide rather than the APR core. The preferential binding of TRA2B-P12 to its homologous tau sites indicates that interaction specificity is encoded in the primary sequence and is not solely a consequence of non-specific hydrophobic attraction.

### Both TRA2B aggregation-prone regions are required for enhanced tau aggregation

The dual-APR architecture of TRA2B is required for function. Peptides encoding APR1 alone (TRA2B-P2) or APR2 alone (TRA2B-P3) had no measurable effect on the aggregation kinetics of Tau297-391, a tau fragment encompassing both PHF6 and PAM4 that aggregates spontaneously without polyanionic cofactors (**Suppl. Fig. 4**). TRA2B-P12, which encodes both motifs in their native sequence context, accelerated tau aggregation in a concentration-dependent manner, shortening the lag phase and shifting the inflection point of the ThT curve (Fig. 4C). TRA2B-P12 alone did not aggregate at the concentrations used, ruling out self-seeding (**Suppl. Fig. 4**). Proline substitution of both APRs in TRA2B-P13 abolished the effect, confirming that the APR sequences rather than the flanking regions are responsible (**Suppl. Fig. 4**).

The same requirement holds for the full-length protein. Proline substitution of both APRs in full-length TRA2B (TRA2B-PP) abolished pro-aggregation activity in tau biosensor cells: cells expressing TRA2B-PP showed tau aggregate formation indistinguishable from GAPDH controls, in contrast to the enhancement produced by wild-type TRA2B (**Fig. 4D-E**). Co-immunoprecipitation established the physical basis for this loss, with tau co-precipitating robustly with wild-type TRA2B but showing markedly reduced association with TRA2B-PP (**Fig. 4F**). Microscale thermophoresis (MST) gave a consistent result: TRA2B-P12 bound tau monomer with an affinity of 4.98 µM, whereas TRA2B-P13 showed very weak binding (**Fig. 4G**). Reduced binding and abolished seeding enhancement therefore track together, indicating that TRA2B promotes tau aggregation through direct, APR-mediated association.

The requirement for both APRs, with two contacts needed and neither sufficient alone, is consistent with the multivalency principle of APR-mediated co-aggregation: simultaneous engagement at the PHF6 and PAM4 interfaces would lower the nucleation barrier by anchoring the incoming molecule to the growing fibril at two points, increasing local avidity. Tau fibrils formed in the presence of TRA2B-P12 showed helical twist that was not observed for tau alone under the same conditions (**Fig. 4H**). This morphology was not present throughout the grid; the predominant species in these samples were short, laterally associated fibrils, and the twisted fibrils showed broad distribution in fibril width and helical pitch (**Suppl fig. 4**). These observations indicate that TRA2B-P12 alters the morphology of tau fibrils formed under these conditions, although the structural basis of this effect remains unknown. If TRA2B and similar proteins can alter tau fibril morphology through APR co-incorporation, they may contribute to the strain diversity of tau pathology observed across tauopathies.

### Full-length TRA2B is a seed-gated switch: dormant in healthy neurons, activated by pathological seeds

The cellular and in vivo data show that TRA2B amplifies tau seeding, yet purified full-length TRA2B has the opposite effect in spontaneous in vitro aggregation assays. At sub-equimolar concentrations, full-length TRA2B inhibited Tau297-391 aggregation, extending the lag phase three- to fivefold and reducing the final aggregated fraction to approximately 30% of the tau-alone control (**Fig. 5A-B**). Full-length TRA2B is a highly positively charged protein, and the disordered regions flanking its RRM may stabilise tau in its soluble form through electrostatic shielding, preventing the APRs from driving spontaneous co-nucleation. Under conditions in which tau is soluble and no seeds are present, TRA2B acts as an inhibitor.

**Figure 5.**
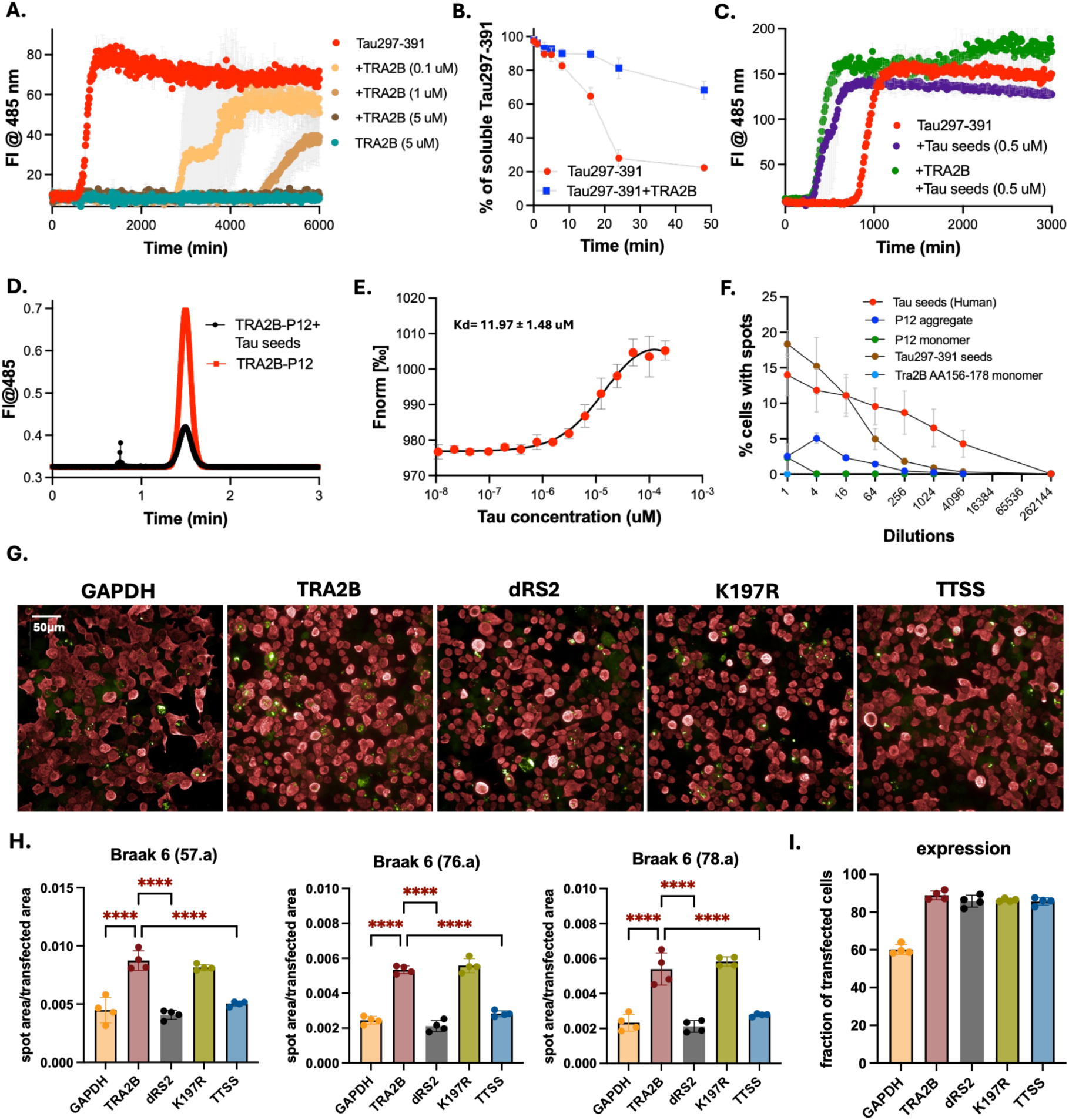
*In vitro* aggregation of Tau and TRA2B. **A**) ThT aggregation kinetics of Tau297–391 in the presence of increasing concentrations of full-length TRA2B (0.1, 1, and 5 μM). **B**) Sedimentation assay showing the loss of soluble Tau297–391 during co-aggregation with TRA2B. **C)** Seeded aggregation of Tau297–391 using Tau seeds in the presence of TRA2B. **D)** Taylorgrams of fluorescently labelled TRA2B-P12 with tau seeds. The reduced amplitude and sharp peaks before the Taylorgram reflect binding of Fluro-P12 to the Tau seeds. **E)** Binding affinity of Fluor-P12 to Tau297-391 seeds (kd=4.98 ± 0.89 uM). The curve is shown as mean values ± SD (n = 4 or 3). **F)** Seeding curves of Tau aggregates following dose-dependent treatment with human Tau seeds (A23162), TRA2B-P12 seeds, TRA2B-P12 monomer, Tau297-391 seeds, or TRA2B AA 156-178 in biosensor cells. **G)** Representative fluorescence images of tau biosensor cells co-transfected with GAPDH (control), wild-type TRA2B, TRA2B lacking the RS2 domain (dRS2), the K197R mutant, or the phosphomimetic variant (TTSS) and with calibrated patient-derived tau seeds from three independent Braak VI cases (57.a, 76.a and 78.a). Scale applies to all images. **H)** Quantification of FRET-positive puncta across all three cases. Deletion of RS2 significantly neutralizes the effect of TRA2B in seeding across all samples. Wild-type TRA2B increased aggregate load in all three cases; both ΔRS2 and TTSS returned it to control levels, whereas K197R did not differ significantly from wild-type TRA2B (n = 4; One-way ANOVA with Tukey’s multiple comparisons test). **I)** Transfection efficiency for GAPDH, wild-type TRA2B, dRS2, the K197R mutant and TTSS variant. Bars show mean ± SD, n = 4

Seeds reverse this behaviour. When preformed tau fibrils were added to Tau297-391 aggregation reactions containing full-length TRA2B, the inhibitory effect was overridden: seeded aggregation proceeded with kinetics at least as fast as seeds alone, regardless of TRA2B (**Fig. 5C**). One possible explanation is that preformed tau fibrils provide a structural template that facilitates engagement of the TRA2B APRs, overcoming the inhibitory influence of the disordered flanking regions.

Flow-induced dispersion analysis (FIDA) supports this: TRA2B-P12 bound tau seeds, whereas TRA2B-P13 did not (**Fig. 5D**). Consistent with this, MST showed that TRA2B-P12 binds tau fibrils (seeds) with an affinity of 11.97 µM (**Fig. 5E**). This seed-gated behaviour resolves the apparent discrepancy between the in vitro and cellular data: TRA2B suppresses spontaneous tau aggregation under the conditions of a healthy neuron, but acts as a co-aggregating amplifier once a seeding template is present. In the diseased brain, where tau seeds are present from early in the pathological process, TRA2B would be constitutively in its amplifying state. This also accounts for the observation that its threshold-lowering effect is largest for the weakest seeds, which require the most assistance to overcome the nucleation barrier.

### TRA2B-derived aggregates cross-seed tau, establishing a self-amplifying cycle

Once TRA2B has been co-incorporated into fibrillar assemblies, as occurs in the insoluble fraction of AD brain, those assemblies acquire the capacity to seed further tau aggregation. TRA2B-P12 fibrils, but not monomers, induced tau aggregate formation in biosensor cells above background, producing FRET-positive puncta in a dose-dependent manner (**Fig. 5F**). The seeding efficiency of TRA2B-P12 fibrils was lower than that of human-derived tau seeds, as expected for a heterotypic rather than homotypic interaction, but cross-seeding was reproducible. This raises the possibility of a positive feedback mechanism: the TRA2B APRs enable co-incorporation into tau aggregates, and those aggregates then become seeding-competent for further tau aggregation, recruiting more tau and potentially more TRA2B. In a neuron in which TRA2B has been displaced from the nucleus by early pathological tau oligomers, such a cycle would progressively amplify both tau burden and TRA2B sequestration.

### The RS2 domain and K197 regulate the tau-amplifying activity of TRA2B

The APRs establish the sequence-specific interaction between TRA2B and tau, but do not explain the striking context dependence of this interaction. TRA2B-P12, which exposes the two APRs in isolation, accelerates tau aggregation in vitro and seeds tau aggregation in cells, whereas full-length TRA2B inhibits spontaneous tau aggregation in vitro but enhances seeded aggregation in cells. This suggested that the APRs are subject to regulation within the full-length protein and that features of TRA2B outside the APR-containing RNA recognition domain may determine whether their interaction with tau becomes productive.

TRA2B has a characteristic SR-protein architecture, with a central RNA-binding domain flanked by arginine/serine-rich regions RS1 and RS2. The C-terminal RS2 domain is particularly rich in potential regulatory sites and contains K197, a lysine that can potentially undergo several post-translational modifications (30). Of particular interest, TRA2B has been reported to undergo SUMOylation, with K197 implicated as a SUMO acceptor site (31). SUMOylation was therefore an attractive candidate mechanism through which the cellular environment might regulate the otherwise cryptic pro-aggregating activity of TRA2B. However, mutation of K197 cannot distinguish SUMOylation from other lysine-dependent modifications, including ubiquitination, acetylation or methylation, and we therefore treat K197 as a regulatory site rather than assigning its phenotype specifically to SUMOylation.

We tested this regulatory layer by comparing wild-type TRA2B with variants that perturb the RS2 domain, K197 and phosphorylation state in tau biosensor cells (**Fig. 5G-I**). Relative to GAPDH-transfected controls, wild-type TRA2B enhanced seeded tau aggregation across all three Braak 6 seed preparations. Deletion of RS2 (TRA2B-dRS2) abolished this enhancement, returning aggregate load to control levels. A phosphomimetic variant in which the RS2 serines/threonines were substituted with glutamate (TRA2B-TTSS) likewise lost the wild-type enhancement, remaining at or marginally above control levels, indicating that phosphorylation mimicry neutralises rather than reverses TRA2B activity. In contrast, mutation of K197 to arginine did not abolish the enhancement. Since dRS2 and TTSS perturb the same regulatory module from two directions while K197R leaves the phenotype largely intact, these data place the requirement for TRA2B-driven enhancement specifically on RS2. Together with the loss of enhancement caused by proline disruption of the TRA2B APRs, these results separate two requirements for cellular tau amplification: intact APRs are required for physical engagement with tau, whereas the RS2 region and K197 determine whether this interaction adopts a pro-aggregating state.

The potential involvement of SUMO is intriguing in light of tau biology (32). Tau SUMOylation at K340 has been reported to promote tau aggregation (33). K340 lies between PHF6 and PAM4, the same two tau APRs recognised by the dual APRs of TRA2B. This raises the possibility that modification of TRA2B at K197 could influence the heterotypic TRA2B–tau interaction more directly—for example, by presenting a modification such as SUMO in trans within an APR-mediated complex. We currently have no evidence that K197 is SUMOylated under our experimental conditions, nor that K197 and tau K340 are spatially juxtaposed, so this remains a hypothesis rather than a mechanistic conclusion. Nevertheless, the combined data identify RS2 and K197 as regulatory elements that control whether the intrinsic tau-binding capacity encoded by the TRA2B APRs is inhibitory, neutral or pro-aggregating.

Intriguingly, TRA2B species consistent with loss of the RS2 domain was also detected in the insoluble fraction of AD brain (**Suppl Fig. 5**). Because experimentally expressed TRA2B-dRS2 suppresses rather than enhances tau aggregation in cells, the presence of this species in human pathological material raises the possibility that RS2 processing accompanies, or counteracts, TRA2B recruitment into insoluble assemblies. Whether the endogenous species arises through proteolytic processing, alternative splicing or another mechanism remains to be established.

### The latent tau-amplifying activity of TRA2B is regulated by protein state and context

These experiments show that TRA2B activity depends on protein state and experimental context. The isolated dual-APR region promotes tau aggregation, whereas full-length TRA2B inhibits spontaneous aggregation in vitro but enhances tau seeding in cells and in vivo. Disruption of the APRs abolishes this enhancement, while perturbation of the regulatory RS2 region shifts TRA2B away from its pro-aggregating state: deletion of RS2 and phosphomimetic substitution are protective, whereas mutation of K197 produces a neutral phenotype. The behaviour of TRA2B therefore cannot be assigned simply to the presence of its tau-compatible APRs; rather, these provide the interaction potential whose functional outcome is modulated by the molecular and cellular context of the full-length protein. Together, these observations define TRA2B as a regulatable tau modifier whose sequence-encoded aggregation potential can be expressed as inhibitory, neutral or amplifying behaviour depending on protein state and experimental context.

## DISCUSSION

Our findings identify TRA2B as a sequence-encoded modifier of tau aggregation whose activity depends on the presence of pathological tau seeds and on the molecular state of TRA2B itself. The underlying mechanism, involving APR homology between TRA2B and tau, may extend to other proteins. The underlying mechanism, involving APR homology between TRA2B and tau, may extend to other proteins. Precedent for such sequence-dependent heterotypic interactions exists across amyloid systems(34, 35). A striking example is the regulation of necroptosis by viral proteins containing RIP homotypic interaction motifs (RHIMs), which form mixed amyloid assemblies with host necroptotic signaling proteins and interfere with functional necrosome formation, thereby suppressing cell death(36). Similar principles have been observed in disease-associated protein aggregation. Medin, which shares sequence homology with amyloid-β (Aβ), can co-aggregate with Aβ and promote its vascular deposition in cerebral amyloid angiopathy(37), as well as modulate its morphology(38), while islet amyloid polypeptide (IAPP), associated with type 2 diabetes, can cross-seed Aβ aggregation(39). Importantly, heterotypic interactions are not restricted to proteins capable of independently forming amyloids. Aggregation-prone mutant p53 can recruit and sequester the related tumor suppressors p63 and p73 through conserved aggregation-prone sequences, compromising their function(40). Similarly, unrelated cellular proteins containing short sequences homologous to Aβ aggregation-prone regions have been shown to enhance Aβ aggregation, with deletion of regions containing these sequences attenuating their effects(41). Together, these examples suggest that local sequence compatibility can enable heterotypic amyloid interactions across distinct proteins, with consequences ranging from enhanced aggregation and functional sequestration to suppression of amyloid-dependent signaling. The human proteome contains thousands of proteins with short sequences homologous to tau APRs. A subset of these, particularly those containing multiple homologous APRs and expressed in vulnerable neuronal populations, may similarly influence tau aggregation depending on their structural context, accessibility and cellular environment.

The property that makes TRA2B a threshold modifier rather than a constitutive promoter of tau aggregation is its seed-gated behaviour. Under unseeded conditions TRA2B is confined to the nucleus, physically separated from cytoplasmic tau and unable to engage in APR-mediated co-nucleation; this is its functional state in a healthy, pre-pathological neuron. The switch to pro-aggregating behaviour requires two concurrent events: the availability of a seeding template, and mislocalisation of TRA2B to the cytoplasm as a consequence of the nuclear envelope disruption that accompanies pathological tau accumulation. Once cytoplasmic, the dual APRs of TRA2B can engage the growing fibril at the structured interface provided by the seed. A functional consequence follows: the contribution of TRA2B to tau seeding is calibrated to the seeding context. Where seeds are potent and abundant, the additional contribution is proportionally modest; where seeds are weak or nascent, as at the earliest stages of spread into a new region, TRA2B engagement may provide the additional nucleation energy needed to cross the threshold. On this model the cell in which TRA2B is nuclear and inactive is protected, whereas the cell in which nuclear envelope disruption has released TRA2B into a seeded cytoplasm is not, and the threshold-lowering activity of TRA2B matters most when tau pathology is establishing itself in a new neuronal population.

The redistribution of TRA2B in AD brain may be more than a passive consequence of aggregation. The shift from the nuclear, TDP-43-correlated compartment to the tau-correlated compartment in late-stage frontal cortex; the biochemical modification of the accumulated TRA2B species; and the neuronal nuclear clearance together indicate that TRA2B is progressively sequestered into tau pathology as disease advances. TRA2B sequestration could have two consequences. First, nuclear depletion of TRA2B disrupts alternative splicing of neuronal transcripts required for synaptic function, neuronal maturation and survival, a broad loss of function. Second, the sequestered protein is itself the regulator of tau pre-mRNA splicing, so loss of nuclear TRA2B activity would be expected to alter the 3R/4R tau isoform balance in affected neurons, potentially shifting isoform composition toward forms with altered aggregation propensity or clearance kinetics. This feedback, in which tau pathology sequesters the splicing factor that regulates tau expression, may contribute to the progressive and self-reinforcing character of tau pathology.

The model we propose is as follows (**Fig. 6**). Under physiological conditions TRA2B is a nuclear splicing factor, and its APRs are embedded in a positively charged, soluble context that prevents co-aggregation with cytoplasmic tau. Early pathological tau oligomers disrupt nuclear envelope integrity, a documented event in tauopathy (42), releasing TRA2B into the cytoplasm, where it encounters seeded tau aggregates. The APRs of TRA2B then engage the growing tau fibril at both the PHF6 and PAM4 interfaces, TRA2B is co-incorporated, and its nuclear pool is depleted. The resulting TRA2B-containing aggregates acquire cross-seeding capacity, amplifying both tau burden and TRA2B sequestration. Splicing regulation of neuronal transcripts, including tau itself, deteriorates as nuclear TRA2B falls below functional levels. These processes could contribute to progressive tau aggregation and TRA2B depletion, further TRA2B loss, isoform imbalance and a progressively lower threshold for seeding of adjacent cells.

**Figure 6.**
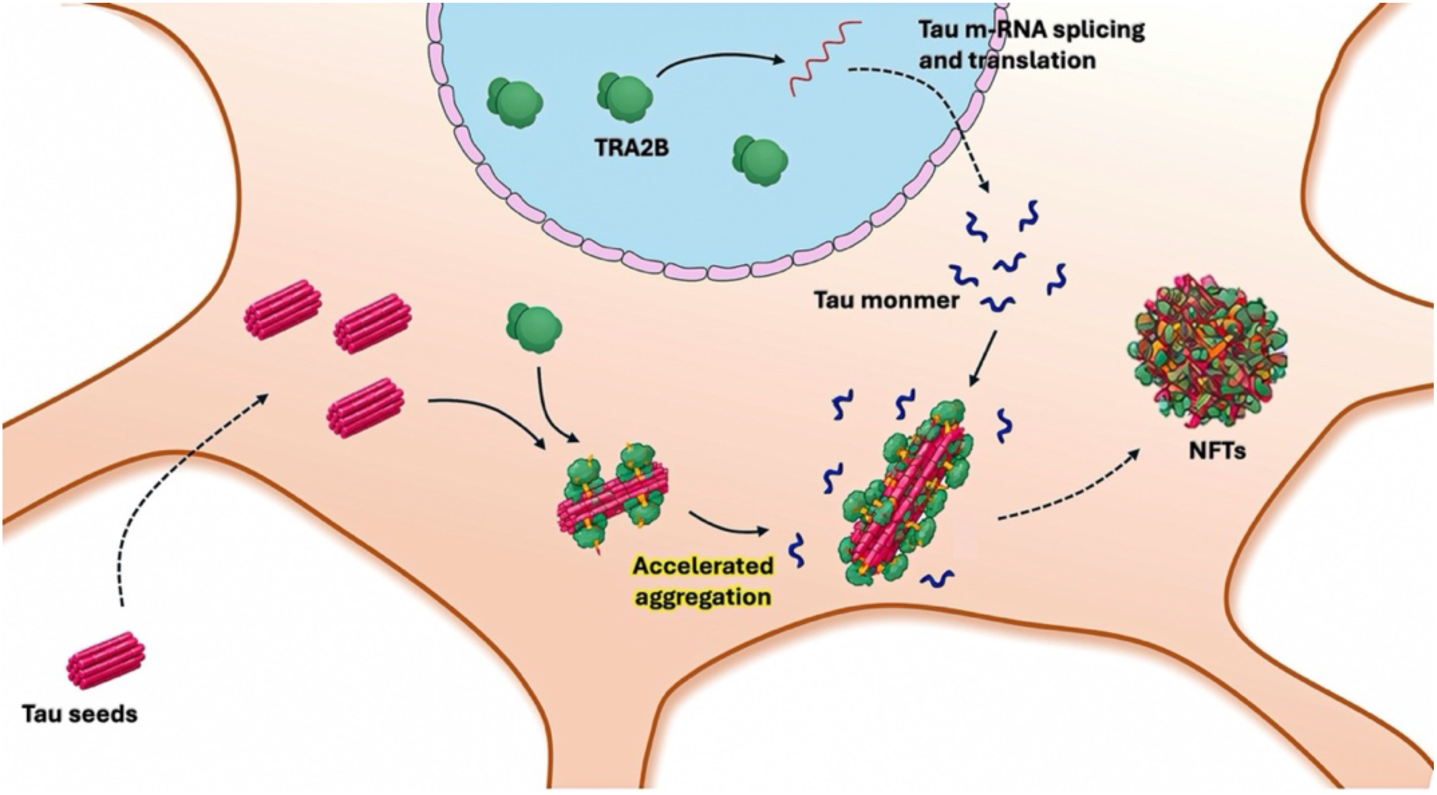
Schematic of TRA2B as a tau amplifier. Cytoplasmic TRA2B engages tau seeds through its APRs, lowering the seeding threshold and amplifying fibril growth. Nuclear TRA2B regulates MAPT splicing and tau translation, generating cytoplasmic tau monomer.

The protective effect of TRA2B-ΔRS2 also raises a potential therapeutic implication. Rather than requiring complete suppression of TRA2B, which would interfere with its physiological function as a splicing factor, it may be possible to selectively uncouple its pathological interaction with tau. Overexpression of TRA2B-dRS2 suppresses tau seeding in cells, suggesting that a TRA2B species lacking the RS2-dependent activation mechanism can oppose rather than promote aggregation. Although the basis of this protective effect remains to be established, it provides proof of principle that the TRA2B–tau interaction may be pharmacologically redirected rather than simply eliminated. Defining how RS2 regulation and the TRA2B APRs cooperate to generate the pro-aggregating state may therefore reveal intervention points for stabilising a non-amplifying, potentially protective state of TRA2B.

Several limitations qualify these conclusions. The human brain data are correlative: the solubility shift, the altered antibody reactivity and the nuclear clearance are each consistent with sequestration of TRA2B into tau pathology, but none demonstrates it directly, and we cannot exclude that TRA2B redistribution is a consequence rather than a cause of aggregation in these tissues. The change in fibril morphology produced by TRA2B-P12 is likewise consistent with co-incorporation but does not establish it, and structural analysis will be required to determine whether TRA2B-derived sequences are part of the fibril core. In vivo, TRA2B was delivered by AAV at levels above the endogenous range, so the magnitude of the effect we measure need not reflect the contribution of endogenous TRA2B; a loss-of-function experiment in the same model would address this directly. Much of the in vitro mechanism rests on Tau297-391 and on short peptides rather than full-length tau, and the affinities we report (4.98 µM for monomer, 11.97 µM for fibrils) are modest, so the cellular relevance of the interaction depends on local concentration and avidity effects that we have not measured. Finally, the nuclear envelope disruption step in the model above is taken from the literature rather than measured in our material, and remains the least constrained element of the proposed mechanism.

These findings also bear on selective neuronal vulnerability in AD. Why specific neuronal populations succumb to tau pathology while adjacent populations are spared has usually been framed in terms of tau-intrinsic properties or clearance mechanisms. Our results suggest that the proteome composition of vulnerable neurons may also contribute, specifically the balance between APR-homologous tau amplifiers and inhibitors that each cell type expresses. A neuron expressing high levels of TRA2B, or enriched for other dual-APR tau co-aggregators, would operate with a lower effective seeding threshold, so that early tau seeds which would stall in a cell with a more protective proteome composition can propagate. As TRA2B and other amplifiers are sequestered into tau pathology, the threshold would fall further and the protective capacity of the nuclear pool would erode. The progressive and region-specific character of tau pathology may therefore reflect, in part, the cell-type-specific composition of this amplifier landscape, which is experimentally accessible through APR homology scanning and whose members are tractable targets for intervention.

## METHODS

### Human autopsy cases

All postmortem brain tissue samples used in this study were obtained from the UZ/KU Leuven Brain Bank, with approval from the ethics committee (Leuven: S59292, S52791, S66705, S63759). Neurofibrillary tangle (NFT) Braak stages were determined on sections immunostained for phosphorylated Tau (p-Tau202/205), following established protocols (43). Neuropathological diagnoses were assigned according to the National Institute on Aging–Alzheimer’s Association criteria(44). Global Clinical Dementia Rating (CDR) scores were retrospectively determined based on published guidelines(45), allowing classification of cases into symptomatic Alzheimer’s disease (AD), asymptomatic AD (asymAD), and healthy non-demented controls (CT). To prevent discrimination, no data on racial background were recorded. All cases were matched for age and sex to the greatest extent possible.

### Mass Spectrometry Sample Preparation

#### 1. Protein Extraction and Fractionation of Human Brain Tissue (Suppl Fig. 6)

Protein extraction was performed on fresh-frozen postmortem human brain tissue (cases 1–78 from Brodmann area 36 [BA36] and 1–80 from BA6). Two BA36 samples were excluded due to insufficient tissue availability; the remaining cases were matched across regions. Approximately 100 mg of grey matter was isolated from each sample, and the leptomeninges and adjacent white matter were carefully dissected away. Tissue was homogenized at 10% (w/v) in cold 1× Tris-buffered saline (TBS) supplemented with protease and phosphatase inhibitors (Thermo Scientific, Waltham, MA, USA) using plastic micropestles (Carl Roth, Karlsruhe, Germany), followed by sonication (6 cycles of 20 s pulse with 10 s rest intervals on ice). To separate four fractions along a solubility gradient, we used a previously established protocol(46). Homogenates were centrifuged at 14,000 × g for 30 min at 4 °C. The supernatant was ultracentrifuged at 175,000 × g for 30 min at 4 °C, yielding pellet II. Pellet II was resuspended in 500 µL 1× TBS containing 1× protease/phosphatase inhibitor and sonicated (1× 20 s) to yield the dispersible (DISP) fraction. The initial insoluble pellet (pellet I) was resuspended in 500 µL TBS containing 2% sodium dodecyl sulfate (SDS; J&K Scientific GmbH, Pforzheim, Germany) and sonicated (5 × 20 s). Following centrifugation (14,000 × g, 30 min, room temperature [RT]), the SDS-soluble supernatant (supernatant III) was collected. Protein fractions were stored at –80 °C until further analysis. Protein concentration was determined using the BCA assay (Pierce MS Grade, Thermo Scientific, Waltham, MA, USA).

#### 2. Protein Precipitation and Sample Concentration

The SDS fraction was precipitated using chloroform/methanol to remove interfering reagents and concentrate proteins. Samples from DISP and SDS fractions were centrifuged (14,000 × g, 20 min, RT) and adjusted to 100 µL with Milli-Q water. Methanol (400 µL), chloroform (100 µL), and water (300 µL) were added sequentially, incubated on ice for 2 min, and centrifuged (14,000 × g, 5 min, 4 °C). The upper aqueous phase was removed, 400 µL methanol was added to the interphase, and samples were centrifuged again (14,000 × g, 5 min, RT). After discarding the supernatant, pellets were washed with 400 µL methanol, centrifuged (14,000 × g, 5 min, RT), and air-dried. Dried pellets were resuspended in 49 µL 50 mM ammonium bicarbonate (ABS; Sigma-Aldrich, Darmstadt, Germany) with 0.02% Rapigest SF (Waters, Milford, MA, USA), incubated overnight at RT with shaking (450 rpm), and stored at –80 °C until tryptic digestion.

#### 3. Tryptic Digestion

Protein concentration was determined using the bicinchoninic acid (BCA) assay (Pierce MS Grade, Thermo Scientific, Waltham, MA, USA). Samples were sonicated in a bath sonicator, and the protein input was normalized to 2 µg in 200 µL of 50 mM ABS. Reduction was performed with 1 mM dithiothreitol (DTT; 45 min, 56 °C; Sigma-Aldrich, Darmstadt, Germany), followed by alkylation with 10 mM iodoacetamide (45 min, room temperature, in the dark; AppliChem). Proteolysis was carried out overnight at 37 °C using MS-grade trypsin (1:100 w/w, enzyme:substrate; Pierce MS Grade, Thermo Scientific, Waltham, MA, USA) with shaking at 300 rpm. Digestion was quenched by acidification with 2 µL trifluoroacetic acid (TFA; Sigma-Aldrich, Darmstadt, Germany) to pH ≈ 2, incubated for 45 min at 37 °C with shaking at 300 rpm, then dried in a SpeedVac and stored at –80 °C until peptide cleanup.

#### 4. Peptide Desalting

Peptides were desalted using C18 StageTips (3M Empore™, C18 Disks, Sigma-Aldrich, Darmstadt, Germany) with acetonitrile and formic acid, dried by SpeedVac, and stored at – 80 °C until LC-MS/MS analysis.

### LC-MS/MS sample acquisition data independent analysis (DIA)

Tryptic peptides were reconstituted in 80 μL of 0.1% formic acid (FA) in LC-MS-grade water, and ∼20 μL were loaded onto Evotips using a Bravo Automated Liquid Handling Platform following Evosep guidelines. Peptides were analyzed on an Evosep One LC system coupled to a ZenoTOF 7600 mass spectrometer (Sciex, Framingham, MA, USA) operated with Sciex OS (v3.3 or higher). Separation was performed using the 60 SPD (samples-per-day) method with a 44-min gradient on an EV1137 Performance column (ReproSil Saphir C18, 1.5 μm; 15 cm × 150 μm) at flow rates of 1–10 μL/min. Mobile phases consisted of 0.1% FA in water and ACN/0.1% FA. The ZenoTOF was operated in SWATH mode with an OptiFlow Turbo V ion source (4500 V spray voltage; gas 1, 20 psi; gas 2, 10 psi; curtain gas, 40 psi; 100 °C). The Zeno SWATH DIA method employed 85 variable-width windows covering 350–1250 m/z. TOF-MS scans were acquired from 350–1250 m/z (25 ms), and TOF-MS/MS scans from 230–1400 m/z (20 ms) using dynamic collision energy, charge state 2, and Zeno pulsing enabled.

### Label-free quantification

All raw data from the ZenoTOF 7600 system (.wiff files) were processed with Spectronaut (version 19) in directDIA+ (deep) mode. Spectra were compared against the Human reference proteome database (https://www.uniprot.org/proteomes/UP000005640) using default Spectronaut settings: precursor and protein Q value (experiment) cutoffs set to 0.01; quantification based on MS2 area; normalization strategy set to “Automatic”; mass tolerance set to default. Oxidation of methionine residues and N-terminus acetylation were considered variable modifications, while carbamidomethylation of cysteine was a fixed modification. The protein quantification reports were exported for further downstream analysis.

### Mass spectrometry data pre-processing

LC-MS/MS data were pre-processed independently for each protein fraction dataset before statistical analysis. Proteins identified as potential contaminants, including keratins (KRT family) commonly introduced during sample handling, were removed. Each dataset was subsequently filtered to exclude proteins with ≥30% missing quantitative values across samples within that dataset. The remaining quantitative values were log₂-transformed, and all subsequent analyses were performed using the filtered, log₂-transformed datasets.

### Extraction of Tau aggregates

Sarkosyl-insoluble material was extracted as previously described (47). Briefly, human or mouse brain tissue was homogenized using a FastPrep (Fisher Scientific, Merelbeke, Belgium) in cold buffer (10 mM Tris-HCl pH 7.4, 0.8 M NaCl, 1 mM EGTA and 10% sucrose) and treated with Universal Nuclease (Pierce, ThermoFisher Scientific, MA, USA) for 30 min at room temperature (RT). Samples were brought to 1% Sarkosyl (Sigma-Aldrich, Darmstadt, Germany), incubated for 1 h with shaking, and centrifuged at 350,000 × g for 1 h at 4 °C. The pellet was washed, recentrifuged, and resuspended in 50 mM Tris-HCl pH 7.4 (100 mg tissue/100 μL).

### Cell seeding

The Tau MTBD P301S FRET biosensor HEK293 cell line (CRL-3275, ATCC-LGC Standards, France) stably expresses Tau-P301S MTBD fused to CFP and YFP, generating a FRET signal upon seeding with tau aggregate (12). Cells were cultured in DMEM with 10% FBS, 1 mM sodium pyruvate, and non-essential amino acids (Gibco, Thermofisher Scientific, MA, USA) at 37 °C and 5% CO₂. For human tau extracts, cells (15,000/well) were plated in poly-D-lysine-coated 96-well PhenoPlates (PerkinElmer, Mechelen, Belgium). After 24 h, cells were co-transfected with plasmid DNA and varying concentrations of brain extract using Lipofectamine 3000 (Thermo Fisher Scientific, MA, USA) according to the manufacturer’s instructions, with minor adaptions for co-transfection with brain extracts. Prior to transfection, tau seeds were sonicated using a Bioruptor Pico (30 s on/30 s off, 10 °C, 15 min; Diagenode, Seraing, Belgium). Forty-eight hours post-transfection, cells were fixed with 4% formaldehyde for 10 min, washed with PBS, permeabilized and blocked, and stained with an anti-HA primary antibody (1:1000, 3724S, Cell Signaling Technology, MA, USA) followed by an Alexa Fluor 647-conjugated secondary antibody (1:1000, ThermoFisher Scientific, MA, USA). Nuclei were counterstained with DAPI (1:5000, ThermoFisher Scientific, MA, USA) for 30 min. After final washes, cellular imaging was performed from multiple independent experiments (n = 3 - 4) using an Operetta CLS high-content analysis system (PerkinElmer, Mechelen, Belgium). Image analysis was performed using the Harmony software and Colombus Plus digital platform (PerkinElmer, Mechelen, Belgium).

For mouse tau extracts, cells (5,000/well) were plated in poly-L-lysine-coated 384-well PhenoPlates (PerkinElmer, Mechelen, Belgium). After 24 h, cells were transfected with mouse tau seeds using Lipofectamine 3000 (Thermo Fisher Scientific, MA, USA) following the manufacturer’s instructions. Prior to transfection, tau seeds were sonicated using a Bioruptor Pico (30 s on/30 s off, 10 °C, 15 min; Diagenode, Seraing, Belgium). Forty-eight hours post-transfection, cells were fixed with 4% formaldehyde for 10 min, washed with PBS, and stained with DAPI (1:1000, Thermofisher Scientific, MA, USA) for 30 min. Following PBS washes, cells were processed for imaging.

### Co-immunoprecipitation

Tau MTBD P301S FRET biosensor cells (150,000/well) were plated in poly-L-lysine-coated 12-well cell culture plates (Greiner Bio-One, Vilvoorde, Belgium) and co-transfected with plasmid DNA and human tau extract as described above. Forty-eight hours post-transfection, cells were washed with cold PBS and lysed in NP-40 lysis buffer supplemented with protease and phosphatase inhibitors. Lysates were incubated on ice for 15 min, and a fraction was retained as input. Co-immunoprecipitation was performed using anti-HA magnetic beads (Thermo Fisher Scientific, MA, USA) according to the manufacturer’s protocol with minor modifications. Cell lysates were incubated with pre-washed beads in the presence of BSA to reduce nonspecific binding for 30 min at RT with rotation. Beads were washed using a modified TBS-T buffer containing 1M urea. Bound proteins were eluted in SDS-PAGE sample buffer by heating at 95°C for 10 min. Input and eluted fractions were analyzed by SDS-PAGE and Western blotting using standard procedures. Proteins were transferred to nitrocellulose membranes, blocked in BSA-containing buffer, and probed with primary (1:1000, 2367S & 2555S, Bioke, Leiden, Netherlands) and secondary antibodies (1:3000, Bio-Rad, CA, USA) before visualization using a ChemiDoc imaging system (Bio-Rad, CA, USA).

### Animal Experiments

All experimental procedures were approved by the local Animal Ethics Committee and comply with international standards for animal welfare (KU Leuven approval: PLL 184/2020). Protocols were authorized by the KU Leuven Institutional Ethics Committee for animal experimentation. Transgenic mice were housed under conditions designed to ensure health and well-being, with controlled temperatures (20–24 °C) and a 12/12 h light-dark cycle to support normal behavior and circadian rhythms. Animals had ad libitum access to water and pelleted rodent chow. To reduce stress-related confounding effects, mice were transferred to the laboratory one week prior to experimental procedures.

### Murine animal model

PS19 transgenic mice express mutant human microtubule-associated protein Tau (MAPT) under the control of the mouse prion protein (Prnp) promoter (29, 48). The transgene carries the P301S mutation and includes four microtubule-binding repeats with one N-terminal insert (4R/1N). Transgenic mice were maintained by continuous backcrossing of heterozygous animals with wild-type C57BL/6 mice. Two-month-old PS19 mice of both sexes received unilateral hippocampal stereotaxic injections (2 µL) of PBS (n = 5) or AAV2/7_CMVie synapsin-intron-HA-GSlinker-hTRA2B (n = 8) and were sacrificed 4 months post-injection. Animals were genotype-confirmed; non-transgenic (WT) animals identified at genotyping were excluded.

### Stereotactic injection and sample collection

PS19 transgenic mice were anesthetized with 2.5% isoflurane and secured in a stereotaxic frame (Stoelting, Wood Dale, IL, USA). Prior to surgery, animals received buprenorphine (0.05 mg/kg; Vetergesic® 0.6 mg/mL, intraperitoneally) for postoperative analgesia and 400 µL of saline subcutaneously to maintain hydration. Stereotaxic injections were performed unilaterally into the hippocampus (ML: 2.0 mm; AP: −2.3 mm; DV: 2.0 mm) using a Hamilton syringe at a rate of 0.1 µL/min, with the needle left in place for 2 minutes after injection. After wound closure and disinfection, mice were allowed to recover in a warm, quiet environment. Following the incubation period, animals were deeply anesthetized with ketamine and medetomidine and transcardially perfused with PBS followed by 4% paraformaldehyde. Brains were harvested, post-fixed in 4% paraformaldehyde for 24 h, embedded in agarose, and sectioned serially using a vibratome (40 μm). Half of the sections were used for biochemical analysis and half for immunohistochemistry (IHC).

### Immunohistochemistry (IHC)

Brain slices spanning the rostro-caudal extent of the hippocampus were selected for immunofluorescent staining. Following the free-floating staining protocol, each slice was washed twice in PBS for 10 min. Slices were then incubated for 1 h at RT in blocking buffer containing 5% normal goat serum and 5% bovine serum albumin (BSA) diluted in PBS. Subsequently, the slices were incubated overnight at 4 °C with the primary antibody solution prepared in blocking buffer, containing rabbit anti-HA (1:500, C29F4, Cell Signaling, Danvers, MA, USA), mouse anti-phospho-tau pS202/T205 (AT8, MN1020, 1:500, Invitrogen, Waltham, MA, USA), and guinea pig anti-NeuN (1:200, 266 004, Synaptic Systems, Göttingen, Germany). After primary antibody incubation, slices were washed twice in PBS and incubated for 1 h at RT with the appropriate secondary antibodies: goat anti-rabbit Alexa Fluor 488 (1:200, Invitrogen, Waltham, MA, USA), goat anti-mouse Alexa Fluor 555 (1:200, Invitrogen, Waltham, MA, USA), and donkey anti-guinea pig Cy5 (1:200, Bio-connect, Huissen, the Netherlands). Following two additional PBS washes, nuclei were counterstained with NucBlue (ThermoFisher, Waltham, MA, USA) for 10 min at RT. Slices were then mounted onto glass slides and coverslipped using freshly prepared Mowiol mounting medium. Imaging was performed using a Nikon Ti2 spinning-disk confocal microscope (CREST Cicero, Genoa, Italy), housed in the Bio-Imaging Core Facility of VIB.

For human brain staining (cases 1, 7, 8, 11, 12, 16, 42, 53, 55, 56, 57, 64, 65, 67, 68, 72, 78, 81, **Table 1**), 5 μm paraffin-embedded sections were deparaffinized and subjected to epitope retrieval in citrate buffer (pH 6) (Agilent-Dako, CA, USA). Endogenous peroxidase activity was blocked (Agilent-Dako), and sections were incubated overnight with rabbit anti-TRA2B_3 antibody (1:50, HPA063597, Sigma-Aldrich, Darmstadt, Germany). Detection was performed using peroxidase-conjugated anti-rabbit EnVision+ Single Reagent (Agilent-Dako) and DAB chromogen (Agilent-Dako), followed by hematoxylin counterstaining and mounting using an Autostainer XL (Leica, IL, USA).

**Table 1.** Human cases used in this study (BA36: Perirhinal cortex / ectorhinal cortex; BA6: Frontal cortex; PMI: Post-mortem interval; CDR: Clinical dementia rating)

| Case | Region | Diagnosis | Age | Sex | PMI | Braak stage_NFT | Aβ_Phase | CDR |
|---|---|---|---|---|---|---|---|---|
| 1 | BA6, BA36 | nAD | 58 | F | 24 | 0 | 0 | 0 |
| 2 | BA6, BA36 | nAD | 74 | F | 24 | 1 | 0 | 0 |
| 3 | BA6, BA36 | nAD | 55 | M | 96 | 0 | 0 | 0 |
| 4 | BA6, BA36 | nAD | 74 | M | 72 | 0 | 0 | 0 |
| 5 | BA6, BA36 | nAD | 73 | M | 48 | 2 | 0 | 0 |
| 6 | BA6, BA36 | nAD | 45 | M | 24 | 1 | 0 | 0 |
| 7 | BA6, BA36 | nAD | 49 | M | 8 | 0 | 0 | 0 |
| 8 | BA6, BA36 | nAD | 46 | M | 29 | 1 | 0 | 0 |
| 9 | BA6, BA36 | nAD | 51 | M | 48 | 0 | 0 | 0 |
| 10 | BA6 | nAD | 39 | F | 4 | 0 | 0 | 0 |
| 11 | BA6, BA36 | nAD | 55 | M | 96 | 1 | 0 | 0 |
| 12 | BA6, BA36 | nAD | 76 | M | 48 | 1 | 0 | 0 |
| 13 | BA6, BA36 | nAD | 28 | M |  | 0 | 0 | 0 |
| 14 | BA6, BA36 | nAD | 68 | F |  | 0 | 0 | 0 |
| 15 | BA6, BA36 | nAD | 66 | M | 12 | 1 | 0 | 0 |
| 16 | BA6, BA36 | nAD | 64 | M | 24 | 1 | 0 | 0 |
| 17 | BA6, BA36 | nAD | 54 | M | 24 | 1 | 0 | 0 |
| 18 | BA6, BA36 | asymAD | 80 | M | 48 | 1 | 4 | 0 |
| 19 | BA6, BA36 | asymAD | 63 | M | 12 | 1 | 2 | 0 |
| 20 | BA6, BA36 | asymAD | 82 | F | 25 | 4 | 5 | 0 |
| 21 | BA6, BA36 | asymAD | 65 | F | 7 | 1 | 1 | 0 |
| 22 | BA6, BA36 | asymAD | 76 | F | 4 | 1 | 3 | 0 |
| 23 | BA6, BA36 | asymAD | 60 | F | 3 | 0 | 2 | 0 |
| 24 | BA6, BA36 | asymAD | 88 | M | 24 | 3 | 1 | 0 |
| 25 | BA6, BA36 | asymAD | 76 | F | 48 | 1 | 0 | 1 |
| 26 | BA6, BA36 | asymAD | 89 | M | 48 | 1 | 4 | 2 |
| 27 | BA6, BA36 | asymAD | 83 | F | 72 | 5 | 5 | 0 |
| 28 | BA6, BA36 | asymAD | 65 | F | 48 | 2 | 1 | 0 |
| 29 | BA6, BA36 | asymAD | 68 | M | 2 | 0 | 3 | 0 |
| 30 | BA6, BA36 | asymAD | 67 | F |  | 2 | 3 | 0 |
| 31 | BA6, BA36 | asymAD | 64 | F | 48 | 0 | 1 | 0 |
| 32 | BA6, BA36 | asymAD | 72 | F | 24 | 2 | 1 | 0 |
| 33 | BA6, BA36 | asymAD | 73 | F |  | 2 | 2 | 0 |
| 34 | BA6, BA36 | asymAD | 78 | F | 72 | 2 | 3 | 0 |
| 35 | BA6, BA36 | asymAD | 77 | F | 48 | 2 | 4 | 0 |
| 36 | BA6, BA36 | asymAD | 89 | M |  | 2 |  | 0 |
| 37 | BA6, BA36 | asymAD | 84 | F | 72 | 2 | 3 | 0 |
| 38 | BA6 | asymAD | 69 | F | 24 | 1 | 1 | 0 |
| 39 | BA6, BA36 | asymAD | 60 | F | 3 | 1 | 4 | 0 |
| 40 | BA6, BA36 | asymAD | 75 | F | 24 | 1 | 2 | 0 |
| 41 | BA6, BA36 | asymAD | 68 | F | 2 | 0 | 3 | 0 |
| 42 | BA6, BA36 | AD | 72 | F | 12 | 4 | 5 | 1 |
| 43 | BA6, BA36 | AD | 78 | F | 72 | 5 | 5 | 2 |
| 44 | BA6, BA36 | AD | 83 | M | 72 | 4 | 5 | 1 |
| 45 | BA6, BA36 | AD | 83 | M | 48 | 2 | 4 | 1 |
| 46 | BA6, BA36 | AD | 69 | M | 36 | 1 | 5 | 3 |
| 47 | BA6, BA36 | AD | 87 | F | 6 | 4 | 5 | 1 |
| 48 | BA6, BA36 | AD | 85 | M | 36 | 3 | 5 | 1 |
| 49 | BA6, BA36 | AD | 81 | F | 72 | 4 | 4 | 1 |
| 50 | BA6, BA36 | AD | 65 | F | 12 | 3 | 4 | 1 |
| 51 | BA6, BA36 | AD | 96 | F | 72 | 4 | 5 | 1 |
| 52 | BA6, BA36 | AD | 79 | F | 48 | 4 | 4 |  |
| 53 | BA6, BA36 | AD | 67 | F | 72 | 5 | 5 | 3 |
| 54 | BA6, BA36 | AD | 69 | F | 24 | 6 | 5 | 3 |
| 55 | BA6, BA36 | AD | 74 | M | 12 | 6 | 5 | 3 |
| 56 | BA6, BA36 | AD | 73 | M | 24 | 6 | 5 | 3 |
| 57 | BA6, BA36 | AD | 79 | F | 24 | 6 | 5 | 3 |
| 58 | BA6, BA36 | AD | 75 | M | 6 | 5 | 5 | 3 |
| 59 | BA6, BA36 | AD | 92 | F | 12 | 5 | 5 | 1 |
| 60 | BA6, BA36 | AD | 86 | F | 24 | 4 | 5 | 2 |
| 61 | BA6, BA36 | AD | 60 | F | 24 | 5 | 5 | 3 |
| 62 | BA6, BA36 | AD | 64 | M | 24 | 6 | 5 | 1 |
| 63 | BA6, BA36 | AD | 87 | M | 24 | 4 | 5 | 2 |
| 64 | BA6, BA36 | AD | 74 | M | 1 | 6 | 5 | 3 |
| 65 | BA6, BA36 | AD | 77 | F | 12 | 5 | 4 | 2 |
| 66 | BA6, BA36 | AD | 62 | M | 24 | 6 | 5 | 1 |
| 67 | BA6, BA36 | AD | 74 | M | 72 | 6 | 5 | 2 |
| 68 | BA6, BA36 | AD | 78 | F | 1 | 5 | 4 | 1 |
| 69 | BA6, BA36 | AD | 71 | M | 12 | 5 | 5 | 3 |
| 70 | BA6, BA36 | AD | 71 | F | 24 | 6 | 5 | 3 |
| 71 | BA6, BA36 | AD | 82 | M | 12 | 2 | 4 | 3 |
| 72 | BA6, BA36 | AD | 65 | M | 12 | 4 | 5 | 3 |
| 73 | BA6, BA36 | AD | 85 | M | 48 | 5 | 5 | 2 |
| 74 | BA6, BA36 | AD | 55 | F | 24 | 6 | 5 | 3 |
| 75 | BA6, BA36 | AD | 86 | M | 9 | 5 | 5 | 2 |
| 76 | BA6, BA36 | AD | 87 | M | 12 | 6 | 5 | 2 |
| 77 | BA6, BA36 | AD | 76 | M | 24 | 5 | 5 | 3 |
| 78 | BA6, BA36 | AD | 57 | M | 12 | 6 | 5 | 3 |
| 79 | BA6, BA36 | AD | 68 | M | 6 | 6 | 5 | 3 |
| 80 | BA6, BA36 | AD | 76 | M | 24 | 5 | 0 | 3 |
| 81 | BA6 | nAD | 27 | M | 48 | 0 | 0 | 0 |

### IHC single-cell quantification, data processing and statistics

Images were acquired using a Nikon spinning-disk confocal microscope. Cells were detected and classified in QuPath (v0.4.4): nuclear, cytoplasmic, membrane and whole-cell compartments were defined and per-compartment mean intensities measured. Neurons were identified by a NeuN classifier; vector-expressing neurons (Vector+) were defined by the classifier from the nuclear vector (Tracer) signal (restricted to the nucleus, reflecting TRA2B’s predominantly nuclear localisation). Only NeuN-positive cells were analysed. Single-cell data were analysed in Python (pandas, NumPy, SciPy, scikit-learn, matplotlib). Sample identifiers were normalised and sections pooled per animal and hemisphere; detections with tau or NeuN intensity below 50 arbitrary units were discarded as debris. The AT8-high threshold was defined as the 95th/99th percentile of whole-cell AT8 intensity among the true-control population (contralateral, Vector− NeuN+ neurons).

The animal was the unit of statistical inference; cells within an animal were treated as technical replicates. For the cell-autonomous test, the AT8-high fraction was computed for Vector+ and Vector− neurons of the injected hemisphere per animal and compared by a paired one-sided Wilcoxon signed-rank test (Vector+ > Vector−; directional by hypothesis and supported by the bulk data), with per-animal odds ratios (Haldane-corrected). Dose-dependence was assessed by (i) binning nuclear vector intensity into deciles and computing the per-animal AT8-high rate per bin (mean ± SEM across animals) and (ii) a per-animal logistic regression of AT8-high status on log10 nuclear vector intensity, testing the slopes against zero across animals. The hemisphere-level comparison (injected vs contralateral) used the same animal-as-unit framework, excluding animals with substantial contralateral vector expression (bilateral transduction). P < 0.05 was considered significant.

### Western blot analysis

For SDS-PAGE and Western blot analysis, human brain samples (cases 15, 2, 8, 7, 11, 42, 71, 78, 65, 72, 57, 76, 68, 67; **Table 1**) were used. Protein concentrations were determined using the Pierce® BCA assay (ThermoFisher Scientific, Waltham, MA, USA). Lysates (60 μg) were separated on Mini-PROTEAN TGX gels and transferred to 0.2 μm nitrocellulose membranes (Bio-Rad, Temse, Belgium). Membranes were blocked and incubated overnight at 4 °C with primary antibodies, including anti-TRA2B_1 /SFRS10 (1:200, ab171082, Abcam, Waltham, MA, USA), anti-TRA2B_2/SFRS10 (1:1000, AB31353, Abcam, Waltham, MA, USA), anti-TRA2B_3 (1:200, HPA063597, Sigma-Aldrich, Darmstadt, Germany), anti-TRA2B_4 (1:200, sc-166829, Santa Cruz, Dallas, Texas, USA) and PHF1 (pS396/pS404; gift from Dr. P. Davies, New York; 1:3000). After washing, membranes were incubated with fluorescent secondary antibodies (Bio-Rad, Temse, Belgium) and imaged using a ChemiDoc MP system. GAPDH conjugated to Rhodamine (Bio-Rad) was used as a loading control.

### ELISA

Total tau levels in brain homogenates were quantified using the Human Total Tau ELISA Kit (KHB0041, Thermo Fisher Scientific, MA, USA) according to the manufacturer’s instructions. Optical density was measured at 492 nm using an ELx808 microplate reader (Agilent-BioTek, CA, USA). Tau concentrations were derived from a standard curve generated in GraphPad Prism (GraphPad Software, Boston, MA, USA), and all samples were assayed in duplicate to ensure accuracy. We similarly quantified TRA2B protein concentration with Human Transformer 2 Beta ELISA Kit (MBS2033687, MyBioSource, San Diego, CA, USA).

### Expression and purification of Tau297-391

The construct was expressed in chemically competent *E. coli* BL21 Star cells. Transformation was carried out using the pET24-His6-SUMO-Tau297-391 plasmid, and the transformed colonies were grown overnight in LB medium supplemented with 50 μg/mL kanamycin at 37°C until the culture reached an OD₆₀₀ of approximately 2.5. This preculture was then diluted to an OD₆₀₀ of 0.05 and incubated again at 37 °C with shaking (150 rpm) until an OD₆₀₀ of 0.5 was achieved. The culture was subsequently shifted to 18 °C for expression induction with 1 mM IPTG and incubated for 15 hours. Cells were collected by centrifugation and resuspended in lysis buffer (50 mM PB, 500 mM NaCl, 15 mM imidazole, pH 7.5) containing 0.3 mM PMSF and protease inhibitors. Lysis was performed by ultrasonication, and the supernatant was separated from cell debris by centrifugation (18,000 rpm, 50 min, 4 °C). The His6-SUMO–Tau297-391 fusion protein was purified from the lysate using Ni-IMAC chromatography on an FPLC system. After loading onto a Ni-NTA column, the resin was washed with buffer A (50 mM PB, 500 mM NaCl, 15 mM imidazole, pH 7.5) and eluted with buffer B (50 mM PB, 500 mM NaCl, 500 mM imidazole, pH 7.5). Elution fractions were assessed by SDS-PAGE to identify those containing the target protein, which were subsequently pooled. DTT was added to a final concentration of 100 mM, and cleavage of the SUMO tag was carried out with ULP1 protease at 4 °C for 14 hours (**Suppl Fig. 7)**. Finally, Tau297-391 was purified via semi-preparative HPLC using a 20–60% acetonitrile gradient in water containing 0.1% TFA over 20 minutes. Collected fractions were analysed by LC-MS, and automated HPLC runs were used for full-scale purification. The pure protein fractions were combined and lyophilized, yielding approximately 4 mg of white powder per liter of culture.

### Expression and purification of TRA2B-FL

The His₆-TRA2B construct was expressed in chemically competent E. coli BL21 Star cells. Transformation was performed using the pET24-His6-TRA2B plasmid, and protein expression and purification were conducted following the same protocol described for Tau297-391. Following Ni-IMAC purification, the collected fractions contained noticeable impurities; therefore, an additional purification step using cation exchange chromatography was performed. The TRA2B-containing fractions from the Ni-IMAC step were pooled and dialyzed overnight against cation exchange buffer. The dialyzed sample was subsequently loaded onto a cation exchange column, and the eluted fractions were analyzed by SDS-PAGE. This step effectively removed most high-molecular-weight contaminants and the majority of low-molecular-weight impurities. However, a minor low-molecular-weight impurity (approximately 10–12 kDa) co-eluted with TRA2B-FL. To eliminate this residual impurity, the pooled protein fractions were further subjected to dialysis using a 10 kDa molecular weight cutoff membrane, resulting in substantially pure TRA2B-FL protein. To validate the identity of the purified TRA2B-FL protein, three different antibodies were evaluated (841: Abcam, ab31353; polyclonal, 957: Sigma-Aldrich, HPA063597; polyclonal, and 958: Abcam, ab171082; monoclonal). These antibodies were tested using both dot blot and western blot analyses (**Suppl Fig. 8)**.

For dot blot analysis, 5 μL TRA2B-FL (20-30 ug) was spotted on the PVDF membrane and incubated in blocking buffer for 60 min at room temperature. The blocking buffer was poured off, and the primary antibody was added for 2 h at room temperature in 1X PBS buffer. The membranes were washed (3 × 10 min) with PBS-Tween 1X (PBST) followed by incubation with secondary goat anti-mouse Alexa680 (1:10000) for 1 h at room temperature. The membranes were washed again three times with PBST, and protein was then detected and developed using an BioRad ChemiDoc MP Imaging System. For western blot analysis, 10 μg of TRA2B-FL was loaded in each well. The standard protocol of Western blot analysis was followed. In both assays, all three antibodies produced clear and specific signals corresponding to the expected molecular weight of TRA2B-FL, indicating successful recognition of the purified protein. These results confirm that the isolated protein is indeed TRA2B-FL and that it retains the epitopes recognized by both polyclonal and monoclonal antibodies.

### Automated synthesis of peptides

Peptide synthesis was conducted automatically using an Intavis MultiPep RSI peptide synthesizer equipped with a vortex agitation system. Reactions were carried out in 5 mL reactor syringes at ambient temperature. Tentagel R Ram resin (loading capacity: 0.20 mmol/g; synthesis scale: 0.025 mmol) was preconditioned by washing in dimethylformamide and dichloromethane (DMF: DCM (50:50), 3 × 1 min). Peptide assembly was performed using the standard Fmoc solid-phase strategy. All Fmoc-protected amino acids and coupling reagents (HBTU and HATU) were dissolved in DMF to obtain 0.5 M stock solutions. N-methylmorpholine (NMM) was dissolved in DMF to yield a 4 M stock solution and 20% piperadine solution was prepared in DMF. Double coupling was employed for each amino acid addition to ensure complete incorporation. In the first coupling step, Fmoc-AA-OH/HATU/NMM was added to the resin, followed by a second coupling using Fmoc-AA-OH/HBTU/NMM. The automated synthesis cycle consisted of three main steps: (a) Fmoc-deprotection, (b) amino acid coupling, and (c) capping. The detailed protocol for synthesis at a 25 µmol scale is described below.

1. Fmoc Deprotection 600 µL of 20% piperidine in DMF, 5 min reaction time 600 µL of 20% piperidine in DMF, 8 min reaction time Wash with DMF (3.6 mL, 7 × 70 s)
2. Amino Acid Coupling Coupling 1: Fmoc-AA-OH/HATU/NMM (5:4.9:10) in DMF, 15 min reaction time Coupling 2: Fmoc-AA-OH/HBTU/NMM (5:4.9:10) in DMF, 15 min reaction time Wash with DMF (3.6 mL, 1 × 70 s) before proceeding to the capping step
3. Capping Step

The capping solution was freshly prepared by sequentially adding 300 µL of 4 M lutidine in DMF and 300 µL of 4 M acetic anhydride (Ac₂O) in DMF directly to the reactor syringe, generating an in situ 2 M capping mixture. The capping reaction was allowed to proceed for 5 minutes, followed by washing with DMF (3.6 mL, 7 × 70 s). Following completion of the synthesis, peptides were cleaved from the resin with TFA/TIS/H₂O (95:2.5:2.5, v/v/v). The cleavage mixture was added dropwise to cold ether, and the precipitated peptide was collected by centrifugation. The pellet was washed twice more with cold ether, then dried. Crude peptides were purified by preparative RP-HPLC and characterised by LC-MS. TRA2B-P12, TRA2B-P13, fluor-TRA2B-P12, and fluor-TRA2B-P13 were synthesized, purified (HPLC), and characterised by LC-MS (**Suppl Fig. 9**).

Peptide arrays were prepared by SPOT synthesis on acid-stable cellulose membranes preloaded with a PEG spacer and a GGS linker (Aims-Scientific), using an Intavis MultiPep RSi automated peptide synthesizer. Peptides were assembled in the C-to-N direction, after which the amino-acid side-chain protecting groups were removed with a cleavage cocktail. Membranes were washed thoroughly with DMF, DCM, and methanol, and stored under vacuum until use. Prior to incubation, membranes were soaked in 50% methanol for 10 min and rinsed three times briefly in PBS-T (PBS, 0.05% Tween-20), then blocked overnight in 1% BSA in PBS-T. Membranes were subsequently washed three times for 5 min in PBS-T, followed by three washes of 5 min in incubation buffer (10 mM phosphate, 150 mM NaCl, 0.05% Tween-20). Fluorescently labelled peptides were diluted in incubation buffer and applied to the membranes for 1 h. After thorough washing in PBS-T, membranes were imaged on a ChemiDoc system (Bio-Rad), and spot intensities were quantified in ImageJ.

### *In vitro* aggregation assay

Tau297-391 stock preparation: Lyophilized Tau297-391 protein was accurately weighed and dissolved in PBS buffer (10 mM phosphate, 150 mM NaCl) to obtain a 200 µM stock solution. The exact concentration was verified by LC-MS by comparing the integrated LC peak area of the sample with that of a standard chromatogram. Any deviation in the calculated concentration was corrected by adjusting the protein amount or dilution accordingly. Peptide stock preparation: Lyophilized peptides were first dissolved in DMSO with vortexing until completely solubilized, followed by dilution with PBS buffer to achieve a final peptide concentration of 1 mM while maintaining 10% (v/v) DMSO in the final stock solution.

Aggregation assay: For aggregation kinetic monitoring, 100 µL of 10 µM Tau297-391solution was dispensed in triplicate into a half-area 96-well clear-bottom fluorescence plate. Thioflavin T (ThT) was added to a final concentration of 20 µM, and each well contained one PTFE bead (diameter: 3/32” ± 0.002”) to promote uniform aggregation. ThT fluorescence was recorded at 37 °C with continuous shaking (500 rpm) using a FLUOstar Omega plate reader (excitation: 440 nm; emission: 480 nm) in 10-minute intervals. The resulting fluorescence data were processed and plotted using GraphPad Prism 8. All the coaggregation and seeding experiments followed the same protocol for measuring the aggregation kinetics.

Transmission electron microscopy (TEM): Five microliters of the sample were applied to glow-discharged TEM grids (30 s) and incubated for 90 s. Grids were rinsed three times with 25 µL Milli-Q water and stained with 2% uranyl formate for 20 s. After air-drying for 15 min, samples were imaged using a JEOL 1400 TEM operated at 80 kV.

### Sedimentation assay

To evaluate the conversion of monomeric Tau297-391 into aggregated form in the presence of TRA2B-FL, sedimentation assay was conducted. Aggregation reactions were prepared in a 96-well plate following the previously described protocol for measuring ThT aggregation kinetics without ThT. Aliquots were collected at defined time points (0, 1, 3, 5, 8, 16, 24, and 48 h) to assess the remaining monomeric protein. For each time point, a 20 µL sample was withdrawn, diluted to 50 µL using MQ water, and centrifuged at 18,000 × g for 15 min at 4 °C to pellet the aggregated Tau297-391. The resulting supernatant, representing the soluble fraction, was analyzed by LC-MS. The LC peak area corresponding to the monomer at t = 0 h was set as 100%, and subsequent changes in peak area were used to calculate the percentage of soluble TRA2B-FL over time.

### Microscale thermophoresis

MST measurements were performed to calculate the binding affinities of TRA2B-P12 with Tau297-391 monomer and fibrils. Fluor-P12 and Fluor-P13 were synthesized via SPPS, with FITC dye introduced at the N-terminus of TRA2B-P12 or TRA2B-P13 through standard amino acid coupling. Fluor-P12 or Fluor-P13 was mixed with and titrated against varying concentrations of Tau297-391 monomer or fibrils in tau buffer (10 mM PBS, 150 mM NaCl). Measurements were recorded on a Monolith NT automated instrument (NanoTemper Technologies GmbH, Germany) using the green laser channel at 5% LED excitation power and medium MST power, at ambient temperature. Affinity constants and data fitting were performed using NanoTemper analysis software (v2.2.4), and results were plotted in Prism.

### Flow-induced dispersion analysis

FIDA was used to examine binding of TRA2B-P12 and TRA2B-P13 to tau fibrils. Measurements were performed on a FIDA1 instrument equipped with a 480 nm excitation source, using a coated capillary. Fibrils assembled from tau297–391 were sonicated prior to use. Fluorescently labelled TRA2B-P12 and TRA2B-P13 (200 nM) were prepared in assay buffer (10 mM phosphate, 150 mM NaCl, 0.05% Pluronic F-127, pH 7.2, 37°C), pre-incubated with sonicated fibrils (2 µM) and injected as the indicator sample; buffer alone was used as the mobile phase and analyte. Data were processed in the FIDAbio software using the automated analysis with baseline correction applied at 1 min.

## DECLARATIONS

### Ethics approval and consent to participate

All postmortem brain tissue used in this study was obtained from the UZ/KU Leuven Brain Bank with approval from the UZ/KU Leuven ethics committee (approval numbers S59292, S52791, S66705 and S63759).

All animal procedures were approved by the Animal Ethics Committee of KU Leuven (approval number PLL 184/2020) and were authorised by the KU Leuven Institutional Ethics Committee for animal experimentation. Procedures comply with international standards for animal welfare.

### Consent for publication

Not applicable. This study contains no individual person’s data in any form that would permit identification.

### Availability of data and materials

All data supporting the findings of this study are available within the article and its supplementary information files, or from the corresponding authors on reasonable request.

### Authors’ contributions

G.T. and R.K. conceived the study, designed and performed the experiments, analysed the data, administered the project, and wrote the original draft. A.E.W., A.G., S.K., G.V.H., F.C., Ladan K., Laleh K., K.M., B.L. and S.O.T. developed methodology, performed experiments and validated results. R.D. critically revised the manuscript and provided substantial input. R.V., C.A.F.A. and D.R.T. provided human brain tissue and neuropathological expertise. N.L. contributed to the conceptualisation of the study. F.R. and J.S. conceived and supervised the study, provided resources, acquired funding and wrote the original draft. All authors reviewed, edited, and approved the final version.

## Acknowledgements

The authors wish to acknowledge the excellent technical assistance of A. Ronisz and S. Ospitalieri. We thank the VIB BioImaging Core at KU Leuven for training, technical support, and access to their instrument parks. We would also like to thank our collaborators from Proteomics Leuven-Laboratory of Applied Mass Spectrometry (LAMaS) for their valuable contributions and support with the proteomics data acquired for this study. Finally, we would like to thank Valentina Zorzini and VIB-KU Leuven Center for Neuroscience Biophysics Technology Unit. We are grateful to the brain donors and their families, without whom this work would not have been possible.

## LITERATURE

1. Kovacs GG. Tauopathies. Handb Clin Neurol. 2017;145:355–68.

2. Braak H, Braak E. Neuropathological stageing of Alzheimer-related changes. Acta Neuropathol. 1991;82(4):239–59.

3. Mudher A, Colin M, Dujardin S, Medina M, Dewachter I, Alavi Naini SM, et al. What is the evidence that tau pathology spreads through prion-like propagation? Acta Neuropathol Commun. 2017;5(1):99.

4. Clavaguera F, Bolmont T, Crowther RA, Abramowski D, Frank S, Probst A, et al. Transmission and spreading of tauopathy in transgenic mouse brain. Nat Cell Biol. 2009;11(7):909–13.

5. Clavaguera F, Akatsu H, Fraser G, Crowther RA, Frank S, Hench J, et al. Brain homogenates from human tauopathies induce tau inclusions in mouse brain. Proc Natl Acad Sci U S A. 2013;110(23):9535–40.

6. DeVos SL, Corjuc BT, Oakley DH, Nobuhara CK, Bannon RN, Chase A, et al. Synaptic Tau Seeding Precedes Tau Pathology in Human Alzheimer’s Disease Brain. Front Neurosci. 2018;12:267.

7. Vogel JW, Iturria-Medina Y, Strandberg OT, Smith R, Levitis E, Evans AC, et al. Spread of pathological tau proteins through communicating neurons in human Alzheimer’s disease. Nat Commun. 2020;11(1):2612.

8. Courade JP, Zetterberg H, Höglinger GU, Dewachter I. The evolving landscape of Alzheimer’s disease therapy: From Aβ to tau. Cell. 2025;188(26):7337–54.

9. Dujardin S, Commins C, Lathuiliere A, Beerepoot P, Fernandes AR, Kamath TV, et al. Tau molecular diversity contributes to clinical heterogeneity in Alzheimer’s disease. Nat Med. 2020;26(8):1256–63.

10. Dujardin S, Bégard S, Caillierez R, Lachaud C, Carrier S, Lieger S, et al. Different tau species lead to heterogeneous tau pathology propagation and misfolding. Acta Neuropathol Commun. 2018;6(1):132.

11. Komarova NL, Thalhauser CJ. High degree of heterogeneity in Alzheimer’s disease progression patterns. PLoS Comput Biol. 2011;7(11):e1002251.

12. Holmes BB, Furman JL, Mahan TE, Yamasaki TR, Mirbaha H, Eades WC, et al. Proteopathic tau seeding predicts tauopathy in vivo. Proc Natl Acad Sci U S A. 2014;111(41):E4376–85.

13. Kavanagh T, Halder A, Drummond E. Tau interactome and RNA binding proteins in neurodegenerative diseases. Mol Neurodegener. 2022;17(1):66.

14. Shapley SM, Shantaraman A, Gadhavi J, Kearney MA, Dammer EB, Duong DM, et al. Proximity Labeling of the Tau Repeat Domain Enriches RNA-Binding Proteins That Are Altered in Alzheimer’s Disease and Related Tauopathies. Mol Cell Proteomics. 2026;25(1):101458.

15. Maziuk BF, Apicco DJ, Cruz AL, Jiang L, Ash PEA, da Rocha EL, et al. RNA binding proteins co-localize with small tau inclusions in tauopathy. Acta Neuropathol Commun. 2018;6(1):71.

16. Lovell MA, Markesbery WR. Ectopic expression of Musashi-1 in Alzheimer disease and Pick disease. J Neuropathol Exp Neurol. 2005;64(8):675–80.

17. Ash PEA, Lei S, Shattuck J, Boudeau S, Carlomagno Y, Medalla M, et al. TIA1 potentiates tau phase separation and promotes generation of toxic oligomeric tau. Proc Natl Acad Sci U S A. 2021;118(9).

18. Wang C, Terrigno M, Li J, Distler T, Pandya NJ, Ebeling M, et al. Increased G3BP2-Tau interaction in tauopathies is a natural defense against Tau aggregation. Neuron. 2023;111(17):2660–74.e9.

19. Meneses A, Koga S, O’Leary J, Dickson DW, Bu G, Zhao N. TDP-43 Pathology in Alzheimer’s Disease. Mol Neurodegener. 2021;16(1):84.

20. Simonetti F, Zhong W, Hutten S, Uliana F, Schifferer M, Rezaei A, et al. Direct interaction between TDP-43 and Tau promotes their co-condensation, while suppressing Tau fibril formation and seeding. Embo j. 2025;44(24):7395–433.

21. Tomé SO, Tsaka G, Ronisz A, Ospitalieri S, Gawor K, Gomes LA, et al. TDP-43 pathology is associated with increased tau burdens and seeding. Mol Neurodegener. 2023;18(1):71.

22. Beerten J, Schymkowitz J, Rousseau F. Aggregation prone regions and gatekeeping residues in protein sequences. Curr Top Med Chem. 2012;12(22):2470–8.

23. von Bergen M, Friedhoff P, Biernat J, Heberle J, Mandelkow EM, Mandelkow E. Assembly of tau protein into Alzheimer paired helical filaments depends on a local sequence motif ((306)VǪIVYK(311)) forming beta structure. Proc Natl Acad Sci U S A. 2000;97(10):5129–34.

24. von Bergen M, Barghorn S, Li L, Marx A, Biernat J, Mandelkow EM, et al. Mutations of tau protein in frontotemporal dementia promote aggregation of paired helical filaments by enhancing local beta-structure. J Biol Chem. 2001;276(51):48165–74.

25. Louros N, Ramakers M, Michiels E, Konstantoulea K, Morelli C, Garcia T, et al. Mapping the sequence specificity of heterotypic amyloid interactions enables the identification of aggregation modifiers. Nat Commun. 2022;13(1):1351.

26. Wang J, Gao ǪS, Wang Y, Lafyatis R, Stamm S, Andreadis A. Tau exon 10, whose missplicing causes frontotemporal dementia, is regulated by an intricate interplay of cis elements and trans factors. J Neurochem. 2004;88(5):1078–90.

27. Drummond E, Pires G, MacMurray C, Askenazi M, Nayak S, Bourdon M, et al. Phosphorylated tau interactome in the human Alzheimer’s disease brain. Brain. 2020;143(9):2803–17.

28. Louros N, Wilkinson M, Tsaka G, Ramakers M, Morelli C, Garcia T, et al. Local structural preferences in shaping tau amyloid polymorphism. Nat Commun. 2024;15(1):1028.

29. Yoshiyama Y, Higuchi M, Zhang B, Huang SM, Iwata N, Saido TC, et al. Synapse loss and microglial activation precede tangles in a P301S tauopathy mouse model. Neuron. 2007;53(3):337–51.

30. Jamros MA, Aubol BE, Keshwani MM, Zhang Z, Stamm S, Adams JA. Intra-domain Cross-talk Regulates Serine-arginine Protein Kinase 1-dependent Phosphorylation and Splicing Function of Transformer 2β1. J Biol Chem. 2015;290(28):17269–81.

31. Hendriks IA, Lyon D, Young C, Jensen LJ, Vertegaal AC, Nielsen ML. Site-specific mapping of the human SUMO proteome reveals co-modification with phosphorylation. Nat Struct Mol Biol. 2017;24(3):325–36.

32. Pozzi B, Mammi P, Bragado L, Giono LE, Srebrow A. When SUMO met splicing. RNA Biol. 2018;15(6):689–95.

33. Luo HB, Xia YY, Shu XJ, Liu ZC, Feng Y, Liu XH, et al. SUMOylation at K340 inhibits tau degradation through deregulating its phosphorylation and ubiquitination. Proc Natl Acad Sci U S A. 2014;111(46):16586–91.

34. Konstantoulea K, Louros N, Rousseau F, Schymkowitz J. Heterotypic interactions in amyloid function and disease. Febs j. 2022;289(8):2025–46.

35. Louros N, Schymkowitz J, Rousseau F. Heterotypic amyloid interactions: Clues to polymorphic bias and selective cellular vulnerability? Curr Opin Struct Biol. 2022;72:176–86.

36. He C, Varghese NR, Keeler EG, Pham CLL, Xie T, Williams B, et al. Structural studies of an antinecroptosis viral:human functional heteroamyloid M45:RIPK3 using SSNMR. Proceedings of the National Academy of Sciences. 2026;123(18):e2426811123.

37. Wagner J, Degenhardt K, Veit M, Louros N, Konstantoulea K, Skodras A, et al. Medin co-aggregates with vascular amyloid-β in Alzheimer’s disease. Nature. 2022;612(7938):123–31.

38. Pradhan B, Kumar ST, Wagner J, Gallardo R, Orlando G, De Vleeschouwer M, et al. Atomic structures of medin and Aβ fibrils reveal polymorphic remodeling in mixed amyloid systems. Nature Communications. 2026;17(1):5804.

39. O’Nuallain B, Williams AD, Westermark P, Wetzel R. Seeding Specificity in Amyloid Growth Induced by Heterologous Fibrils*. Journal of Biological Chemistry. 2004;279(17):17490–9.

40. Xu J, Reumers J, Couceiro JR, De Smet F, Gallardo R, Rudyak S, et al. Gain of function of mutant p53 by coaggregation with multiple tumor suppressors. Nature Chemical Biology. 2011;7(5):285–95.

41. Konstantoulea K, Guerreiro P, Ramakers M, Louros N, Aubrey LD, Houben B, et al. Heterotypic Amyloid β interactions facilitate amyloid assembly and modify amyloid structure. Embo j. 2022;41(2):e108591.

42. Yuan S, Essepian N, Roberts R, Sherman E, Wang Ǫ, Erisir A, et al. Tau oligomerization induces nuclear lamina invagination and chromatin remodeling in Alzheimer’s disease. Acta Neuropathol. 2026;151(1).

43. Braak H, Alafuzoff I, Arzberger T, Kretzschmar H, Del Tredici K. Staging of Alzheimer disease-associated neurofibrillary pathology using paraffin sections and immunocytochemistry. Acta Neuropathol. 2006;112(4):389–404.

44. Hyman BT, Phelps CH, Beach TG, Bigio EH, Cairns NJ, Carrillo MC, et al. National Institute on Aging-Alzheimer’s Association guidelines for the neuropathologic assessment of Alzheimer’s disease. Alzheimers Dement. 2012;8(1):1–13.

45. Morris JC. The Clinical Dementia Rating (CDR): current version and scoring rules. Neurology. 1993;43(11):2412–4.

46. Li X, Tsolis KC, Koper MJ, Ronisz A, Ospitalieri S, von Arnim CAF, et al. Sequence of proteome profiles in preclinical and symptomatic Alzheimer’s disease. Alzheimers Dement. 2021;17(6):946–58.

47. Goedert M, Spillantini MG, Cairns NJ, Crowther RA. Tau proteins of Alzheimer paired helical filaments: abnormal phosphorylation of all six brain isoforms. Neuron. 1992;8(1):159–68.

48. Takeuchi H, Iba M, Inoue H, Higuchi M, Takao K, Tsukita K, et al. P301S mutant human tau transgenic mice manifest early symptoms of human tauopathies with dementia and altered sensorimotor gating. PLoS One. 2011;6(6):e21050.

