## Supplemental information for "A latent tau amplifier in the neuronal proteome lowers the seeding threshold for Alzheimer’s disease pathology"

### SUPPLEMENTARY INFORMATION

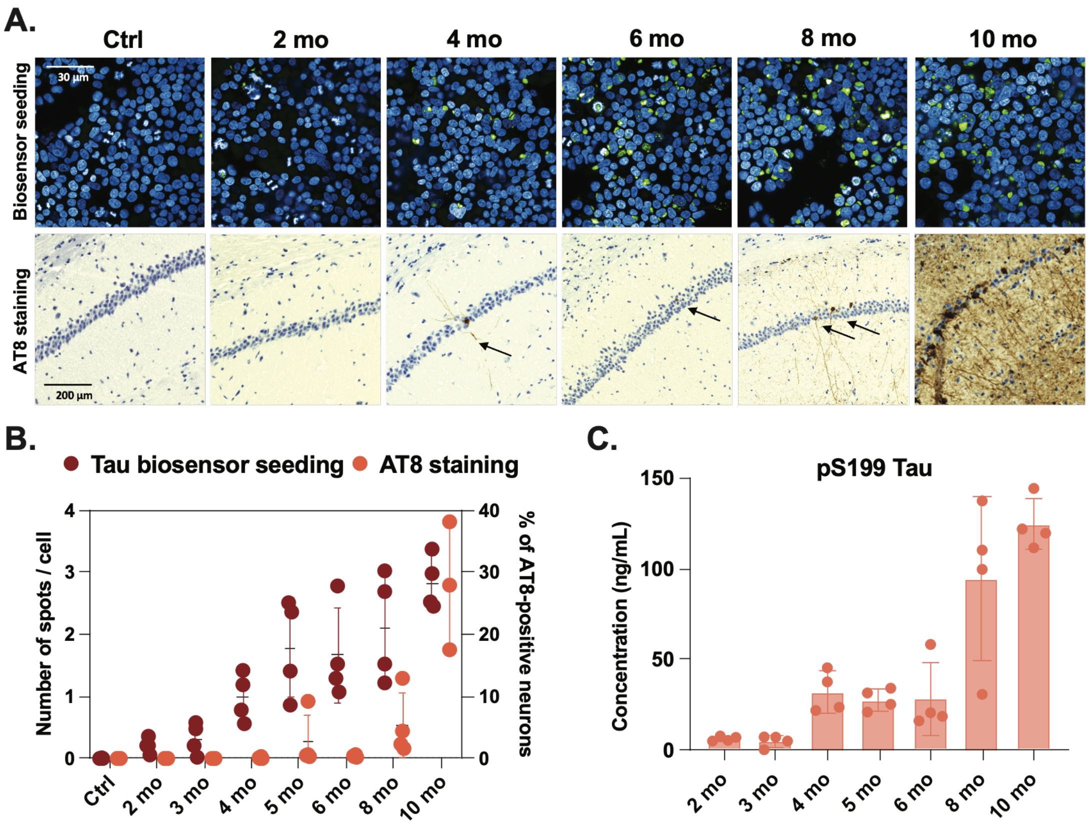


**Supplementary figure 1** Longitudinal characterization of tau pathology in PS19 mice and inhibition of PS19-derived seeding by ScFv-16B12. A) Multi-modal analysis of tau pathology progression in PS19 transgenic mice from 2 to 10 months of age, with representative images of tau seeding activity (FRET-based biosensor assay using sarkosyl-insoluble fractions) and IHC detection of phosphorylated tau (pS203/T205). B) Temporal progression of tau seeding competence and histopathological changes. Seeding activity was detectable as early as 2 months of age, preceding both biochemical and histological markers. A marked escalation in pretangle and neurofibrillary tangle pathology occurred after 8 months (n = 4). C) Quantitative assessment of phosphorylated tau species by ELISA across the longitudinal time course. Biochemical detection of pathological pS199-tau became apparent at approximately 4 months, while IHC-based detection was first observed at 5 months (n = 4).

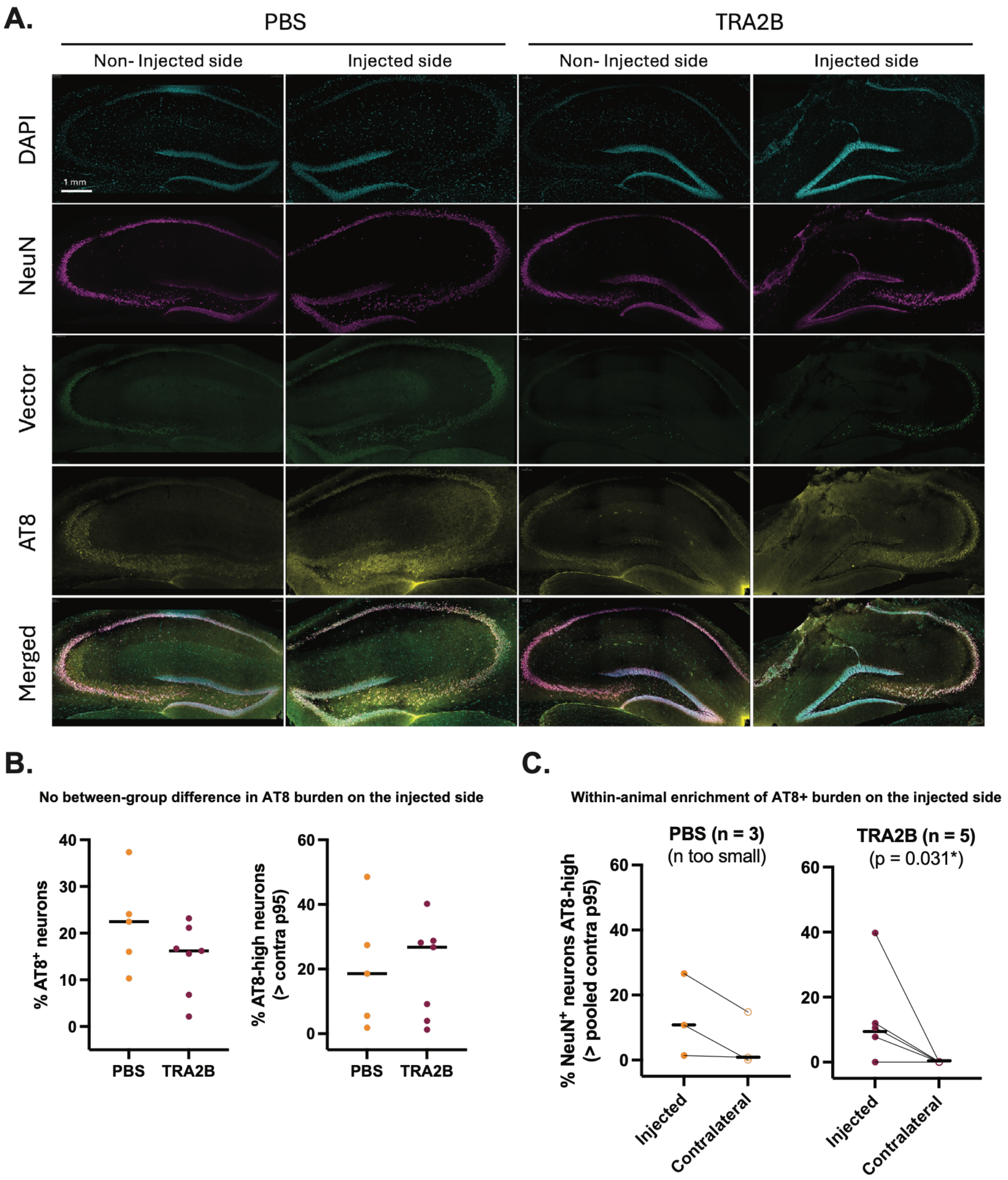


**Supplementary figure 2** Hemisphere-level comparison of AT8-positive neuronal burden in vehicle (PBS)- or AAV-TRA2B- injected animals. **A)** Representative images of hippocampus from animals receiving a unilateral intrahippocampal injection of PBS or AAV-TRA2B and immunostained for DAPI (nuclei), NeuN (neurons), HA (TRA2B vector expression), and AT8 (pS202/T205 tau). Scale bar applies to all images. **B)** Between-group comparison of AT8 burden on the injected hemisphere. Left, percentage of neurons scored AT8-positive; right, percentage of AT8-high neurons, defined as those exceeding the pooled contralateral 95th-percentile AT8 gate. Each point is one animal (PBS, n = 5; TRA2B, n = 7) **C)** Percentage of NeuN$^+$ neurons exceeding the pooled contralateral 95$^{th}$-percentile AT8 gate, injected versus contralateral, per animal; bars, medians; P, one-sided Wilcoxon (injected > contralateral). This comparison is conservative owing to contralateral vector expression. The TRA2B panels are reused from the corresponding panels in Figure 3E.

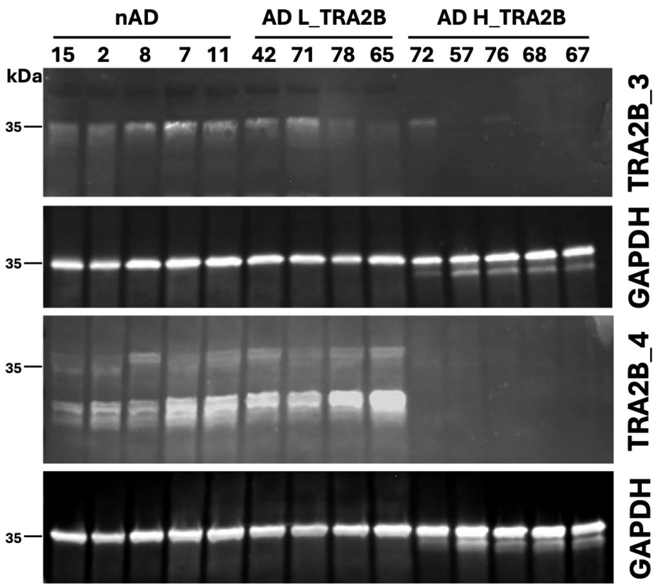


**Supplementary figure 3** Immunoblot of the SDS fraction with anti-TRA2B antibodies. TRA2B is not detected in AD H_TRA2B cases. GAPDH serves as loading control.

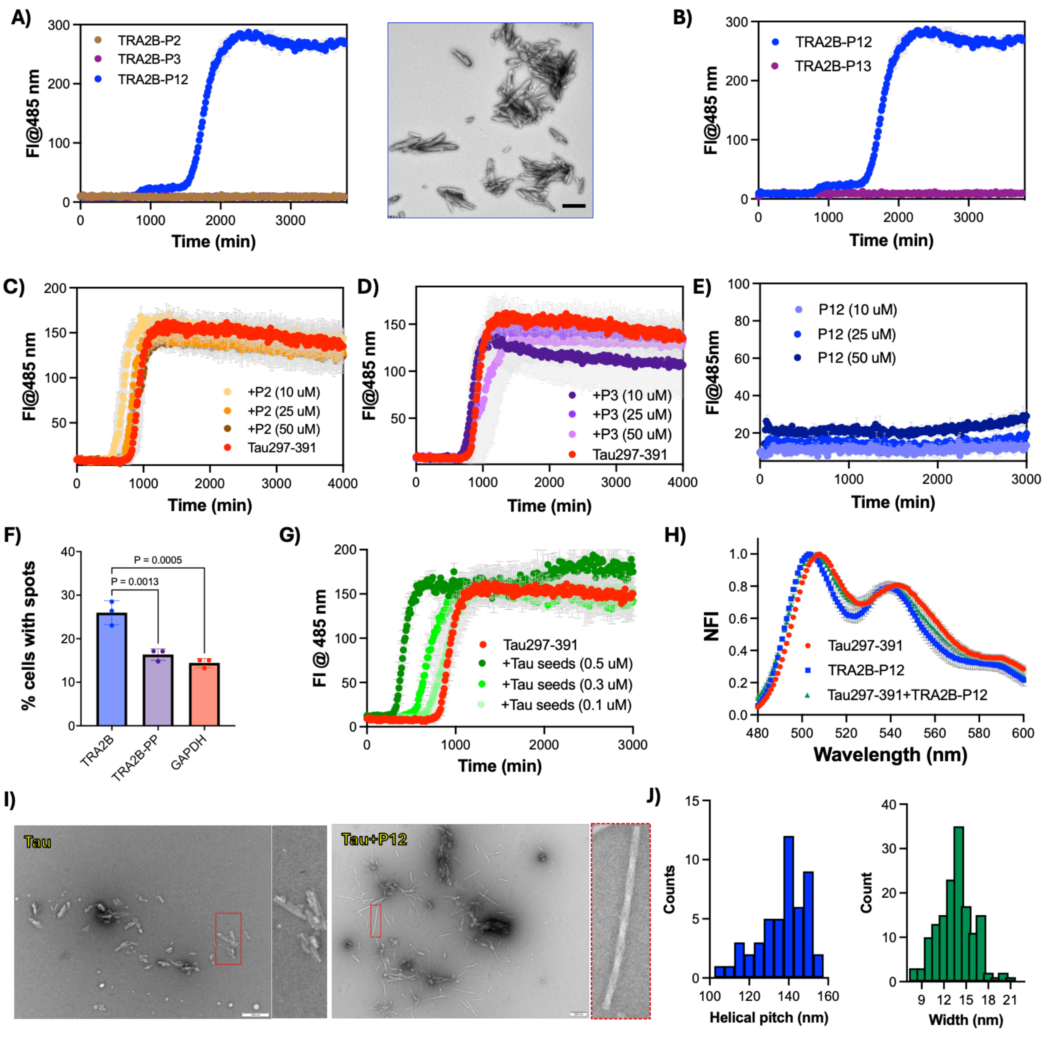


**Supplementary figure 4** Aggregation kinetics measured by ThT fluorescence at 485 nm (mean ± standard error of the mean [SEM], n = 3) for **A)** TRA2B-P2, TRA2B-P3, and TRA2B-P12 at 300 uM and EM images of the fibrils formed by TRA2B-P12. **B)** Aggregation kinetics of TRA2B-P12 (300 uM) and TRA2B-P13. Tau297-391 in the presence of **C)** TRA2B-P2, and **D)** TRA2B-P3. **E)** Aggregation kinetics of TRA2B-P12 at different concentrations. **F)** Seeding of Tau with human Tau seeds in biosensor cells overexpressing wild-type TRA2B, proline-substituted mutant TRA2B-PP, or GAPDH (control). Bar plot (right) highlights differences observed at the highest seed concentration. Data are represented as mean values ± SD (n = 3 independent experiments). **G)** Seeded aggregation of Tau297-391 using increasing concentrations of Tau seeds (0.1, 0.3, and 0.5 µM). **H)** Normalised binding spectra of pFTAA from fibrils derived from Tau297-391, TRA2B-P12, and Tau297-391+TRA2B-P12. Data points are shown as mean values ± SD. **I)** TEM image of fibrillar aggregates from Tau297-391 and Tau297-391+TRA2B-P12 after 72 h. A magnified view of the fibril morphology is shown on the right side of each panel. **J)** Distributions of fibril width (13.7 ± 2.4 nm, n = 150) and helical pitch (137 ± 12 nm, n = 49), measured from negative-stain EM of Tau297–391 co-aggregated with TRA2B-P12.

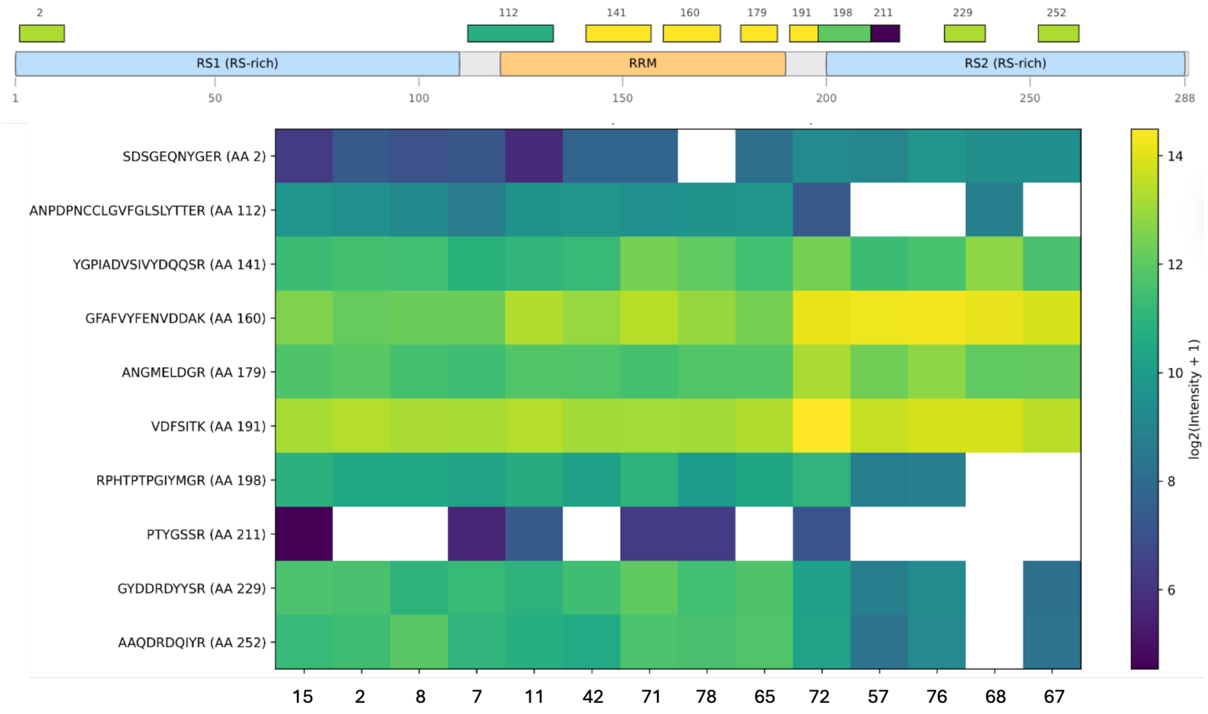


**Supplementary figure 5** RS2 loss in the insoluble fraction of AD brain**.** Position of the identified TRA2B peptides on the domain map of TRA2B, labelled by start residue. The 10 peptides cover 118 of 288 residues (41%). Heatmap of the abundance of these peptides, ordered N- to C-terminally by start residue, in controls (15, 2 8, 7, 11), AD with low TRA2B (42, 71, 78, 65), and AD with high TRA2B (72, 57, 76, 68, 67). Values are log2(intensity + 1), white cells denote peptides not detected in that sample.

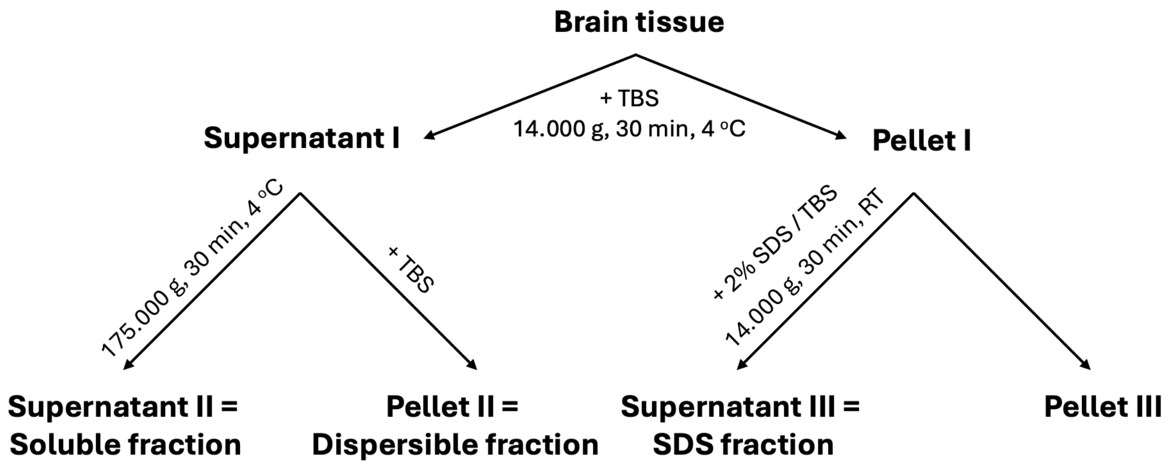


**Supplementary figure 6** Schematic representation of the stepwise biochemical fractionation protocol applied to post-mortem human brain samples for the separation of soluble and insoluble protein fractions.

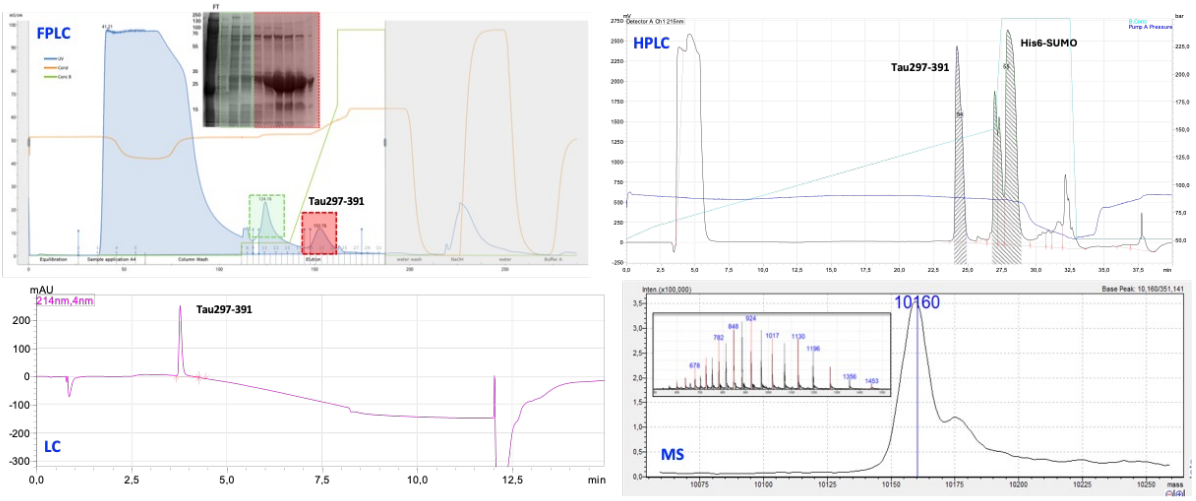


**Supplementary figure 7** Expression and purification of Tau297-391. Representative chromatogram of the Ni-IMAC purification (Inset: analysis by SDS-PAGE of the purification fractions), RP-HPLC chromatogram for the purification and characterization by LC-MS.

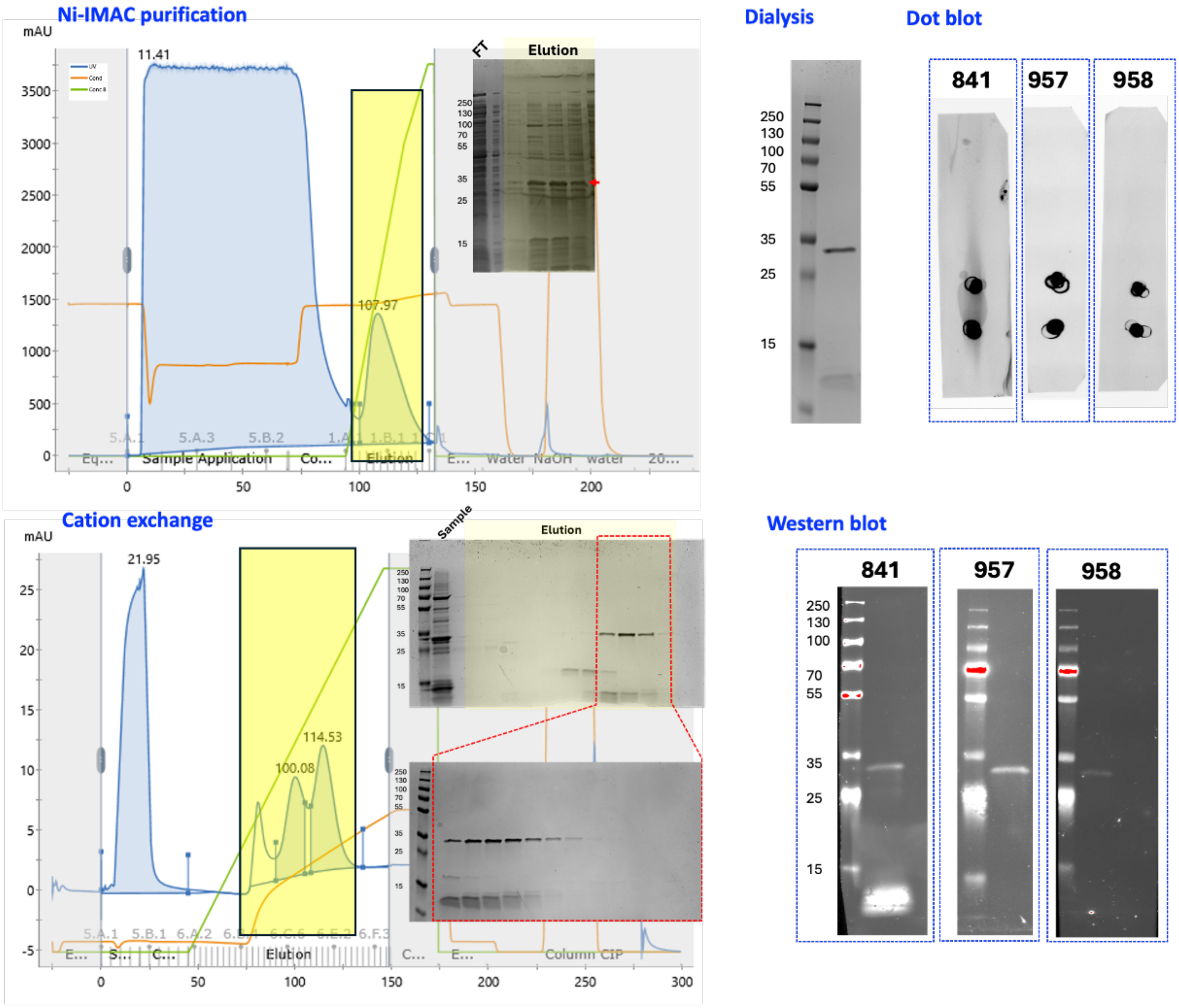


**Supplementary figure 8** Expression and purification of TRA2B-FL. Representative chromatogram of the Ni-IMAC purification (Inset: analysis by SDS-PAGE of the purification fractions), cation exchange chromatogram (Inset: analysis by SDS-PAGE of the purification fractions) for the purification, and Dot blot and western blot analysis of TRA2B-FL.

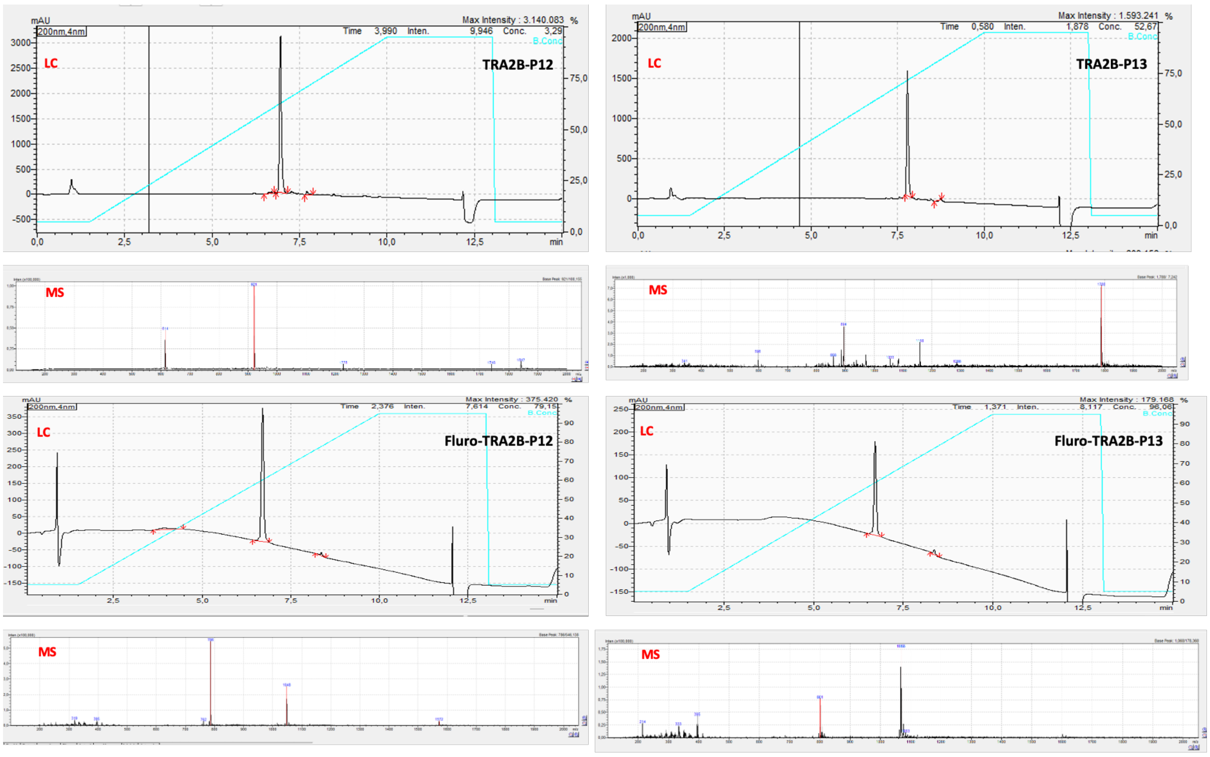


**Supplementary figure 9** LC-MS analysis of TRA2B-P12, TRA2B-P13, fluor-TRA2B-P12, and fluor-TRA2B-P13.
